# Repeated evolution of reduced visual investment at the onset of ecological speciation in high-altitude *Heliconius* butterflies

**DOI:** 10.1101/2024.06.12.598660

**Authors:** David F Rivas-Sánchez, Camilo Salazar, Carolina Pardo-Diaz, Richard M Merrill, Stephen H Montgomery

## Abstract

Colonisation of new habitats is typically followed by divergent selection acting on traits that are immediately important for fitness in the new habitat. For example, shifting sensory environments are often associated with variation in sensory traits critical for navigation and foraging. However, the extent to which the initial response to novel sensory conditions is mediated by phenotypic plasticity, and its contribution to early species divergence remains unclear. We took advantage of repeated cases of speciation in *Heliconius* butterflies with independent allopatric distributions in the west of the Colombian and Ecuadorian Andes. Using volumetric brain measurements, we analysed patterns of investment in sensory processing in brain components across different localities and habitats. We find that a higher-altitude species, *H. chestertonii*, differs in levels of investment in visual and olfactory brain centres compared to its lower altitude relative *H. erato venus*, mainly attributable to heritable variation as inferred from comparisons between wild and common-garden reared individuals. We compared these shifts with those reported for another high-altitude species, *H. himera*, and its parapatric lowland counterpart, *H. erato cyrbia*, and demonstrate parallel reductions in the size of specific optic lobe neuropils. Conversely, for the antennal lobe, we detected disparate trait shifts in *H. himera* and *H. chestertonii* in respect to their lowland *erato* neighbours. Overall, our findings add weight to the adaptive potential for neuroanatomical divergence related to sensory processing during early species formation.

**Lay summary:** Repeated associations between trait variation and environmental shifts may indicate adaptation to local sources of natural selection. For instance, in fish, the presence of certain morphological traits in specific ecological conditions across independent populations is well documented, suggesting equivalent phenotypic responses to shared sources of natural selection. We compared independent cases of ecological divergence in *Heliconius* butterflies distributed along altitude gradients from sea level to mid mountain in the west of the Colombian and Ecuadorian Andes. Shifts in altitude involve repeated, abrupt transitions from wet, large-leaved, warm forests to higher dry, open, cold scrubs. We tested hypotheses about the role of these ecological shifts in driving adaptive evolution in neuroanatomical traits during early speciation. We showed that in *Heliconius*, independent changes in forest-type have been accompanied by heritable parallel patterns of divergence in sensory investment in visual processing in the brain. We propose these differences likely facilitate species divergence in the face of ongoing geneflow.

## Introduction

Local adaptation may promote the evolution of reproductive isolation between populations exposed to different environments due to selection against maladapted migrants or hybrid offspring (Coyne & Orr, 2004; Nosil et al., 2005; Rundle & Nosil, 2005; Schluter, 2009).Traits important for habitat use and resource acquisition are likely exposed to strong divergent selection following habitat colonisation (Via, 2009). In particular, because navigation and foraging require the brain to constantly interpret and respond to external cues, selection on the sensory and neural traits that underlie perception and behaviour is expected to track changes in habitat properties such as food distribution or light-regime (Dell’Aglio et al., 2024; Wcislo, 1989). Classic examples of this process largely stem from aquatic systems. For example, the brain structure of (blind) Mexican cave fish (*Astyanax mexicanus*) differs from that of their surface-dwelling counterparts, (Loomis et al., 2019), and divergence in the visual system of sympatric *Pundamilia* cichlids contributes to lower fitness when reciprocally transplanted to non-native light environments (Maan et al., 2017).

Which traits facilitate species divergence remains an open question, but the relative contribution of selection to phenotypic evolution is often easier to discern between closely related taxa (Gonda et al., 2013). In particular, populations living in similar ecological conditions, and which have evolved similar trait-shifts, likely reflect shared sources of selection, providing evidence of adaptive divergence as opposed to neutral drift (Langerhans & DeWitt, 2004; Losos et al., 1998; Schluter, 2000). For example, freshwater sticklebacks (*Gasterosteus aculeatus*) have independently lost armour plates relative to their marine conspecifics (Colosimo et al., 2005), and parallel divergence in hostplant use has been reported in stick insects (*Timema cristinae*) (Nosil et al., 2002). Both cases suggest habitat-shifts synchronised with divergent adaptations that arise repeatedly under similar selection regimes.

Butterflies occur across a broad range of ecological conditions and provide an excellent opportunity to explore how the sensory and cognitive requirements of contrasting habitats are met by adaptations in peripheral and higher-order brain structures (Couto et al., 2020). Here, we use closely related species from within the Neotropical *Heliconius erato* complex to explore how neuroanatomical trait-shifts may be associated with changes in habitat. Within this group, *H. chestertonii* and *H. himera* (species *sensu* Lamas and Jiggins (2017)) are notable for their specialisation to higher elevation forests in the west of the Colombian and Ecuadorian Andes, respectively (Arias et al., 2008; Jiggins et al., 1996). At their low altitudinal boundaries, these species are in contact with lowland populations of *H. erato*, with which they form narrow hybrid zones (*H. chestertonii* with *H. erato venus*, and *H. himera* with *H. erato cyrbia, H. e. favorinus* and *H. e. emma*) (Arias et al., 2008; Jiggins et al., 1996). In each case, the proportion of hybrids is below that expected under random mating, despite only limited hybrid inviability (McMillan et al., 1997; Muñoz et al., 2010). This is consistent with persistent gene-flow countered by the combined effects of assortative mating and frequency-dependent selection on warning colour patterns (Mallet et al., 1998; McMillan et al., 1997; Merrill et al., 2014; Muñoz et al., 2010). Selection is additionally expected to relate to abrupt environmental shifts between the dense, warm, wet, and climatically stable forests inhabited by lowland *H. erato* populations, and the open, drier and more climatically fluctuating high elevation forests where *H. chestertonii* and *H. himera* are distributed (Arias et al., 2008; Jiggins et al., 1996).

Lower elevation *H. e. cyrbia* and higher elevation *H. himera*, have diverged in traits related to physiology, life-history, development, and flight behaviour (Davison et al., 1999; Dell’Aglio et al., 2022). This is closely mirrored in the ecologically equivalent species pair *H. e. venus*-*H. chestertonii*, implying independent, parallel adaptations to high-altitude forest (Rivas-Sánchez, Gantiva-Q, et al., 2023; Rivas-Sánchez, Melo-Flórez, et al., 2023). *H. e. cyrbia* and *H. himera* also exhibit distinct neuroanatomies that result in greater relative tissue investment in the optic lobe of *H. e. cyrbia* and the antennal lobe of *H. himera* (Montgomery & Merrill, 2017). These differences are heritable and have been interpreted as divergent adaptations to meet the sensory conditions of their respective forest environments (Montgomery & Merrill, 2017). This conclusion is supported by behavioural data showing that contrasting sensory investment predicts differential weighting of information from these sensory modalities during foraging (Dell’Aglio et al., 2022). Additional evidence of the adaptive significance of these shifts could be obtained if they are replicated in a separate, ecologically similar divergence event.

We predicted that *H. chestertonii*, a species genetically independent (Van Belleghem et al., 2021) but ecologically equivalent to *H. himera* (Rivas-Sánchez, Melo-Flórez, et al., 2023), would exhibit neuroanatomical trait-shifts relative to lowland *H. e. venus*, that are similar to those observed between *H. e. cyrbia* and *H. himera*. This is expected if divergent ecological selection along altitudinal gradients from wet to dry forests is met with brain structure adaptations in these butterflies. To test this, we drew comparisons using volumetric data obtained from wild-caught and insectary-reared *H. e. venus* and *H. e. chestertonii*. We then combined our data with previously published brain structure measurements of *H. e. cyrbia* and *H. himera* (Montgomery & Merrill, 2017) to formally test for parallel neural evolution.

## Methods

### i) Collections in the field and in common garden

We collected wild *H. e. venus* (n=20, males= 13, females=7) and *H. chestertonii* (n=18, males=12, females=6) using hand nets in forests located in southwest Colombia at low (Buenaventura, N03º50’0.04” W77º15’45.1” and La Barra, N03°57.558420 W 77°22.705080) and high altitude (Montañitas N03º40’36.3” W76º31’19.4” and El Saladito N03º29’11.5” W76º36’39.8”) in January 2020 and January 2021. Because these sites are distant from the contact zone, hybrids have never been reported in these areas, thus individuals can be considered “faithful” representatives of their species. To explore potential habitat effects on brain structure, we also included first generation insectary-reared *H. e. venus* (n=17, males=4, females=13) and *H. chestertonii* (n=17, males=7, females=10) bred from independent wild-caught females. These butterflies were reared in the José Celestino Mutis research station in La Vega (1485 m), Colombia and sampled after they reached maturity (∼10 days), when post-eclosion brain-growth in *Heliconius* plateaus (Montgomery, Merrill, et al., 2016). To test for parallel trait-shifts in high-altitude populations, we additionally incorporated volumetric brain measurements of wild-caught *H. e. cyrbia* (n=16, males=8, females=8) and *H. himera* (n=16, males=8, females=8), obtained from Montgomery and Merrill (2017).

### ii) Brain tissue preparation

We dissected, fixed and prepared whole brains for indirect immunofluorescence staining against synapsin, a protein expressed in presynaptic sites (anti-SYNORF1; Developmental Studies Hybridoma Bank, Department of Biological Sciences, University of Iowa, Iowa City, IA; RRID: AB_2315424), with a Cy2 conjugated affinity-purified polyclonal goat antimouse IgG (H+L) secondary antibody (Stratech Scientific, Newmarket, Suffolk, UK; Jackson ImmunoResearch Cat No. 115-225-146, RRID: AB_2307343). This protocol reveals borders between neuropils (synapse dense brain regions) and whole brain 3D structure (Ott, 2008).

### iii) Confocal microscope imaging and volumetric reconstructions

We performed imaging on a confocal laser-scanning microscope (Leica TCS SP5 or SP8; Leica Microsystem) with a 10× dry objective (numerical aperture 0.4) to produce 512 × 512 pixels x-y resolution image stacks with a 2 μm z-step. We used Amira 5.5 (Thermo-Fisher Scientific) to merge these image stacks and assign image regions to neuropils of interest. This allowed us to reconstruct and obtain volumetric measurements of sensory neuropils in the optic lobe (lamina, medulla, lobula plate, lobula, ventral lobula, lobula and accessory medulla) and the central brain (antennal lobes, anterior (AOTu) and posterior optic tubercles (POTu)). We measured the total volume of the central brain and subtracted the volume of the antennal lobes, the AOTu, the POTu, the mushroom bodies, the protocerebral bridges and central complex (CX) to obtain the volume of the undifferentiated regions of the central brain (rCBR), which was used as an allometric control in our analyses (Figure 1).

**Figure 1.**
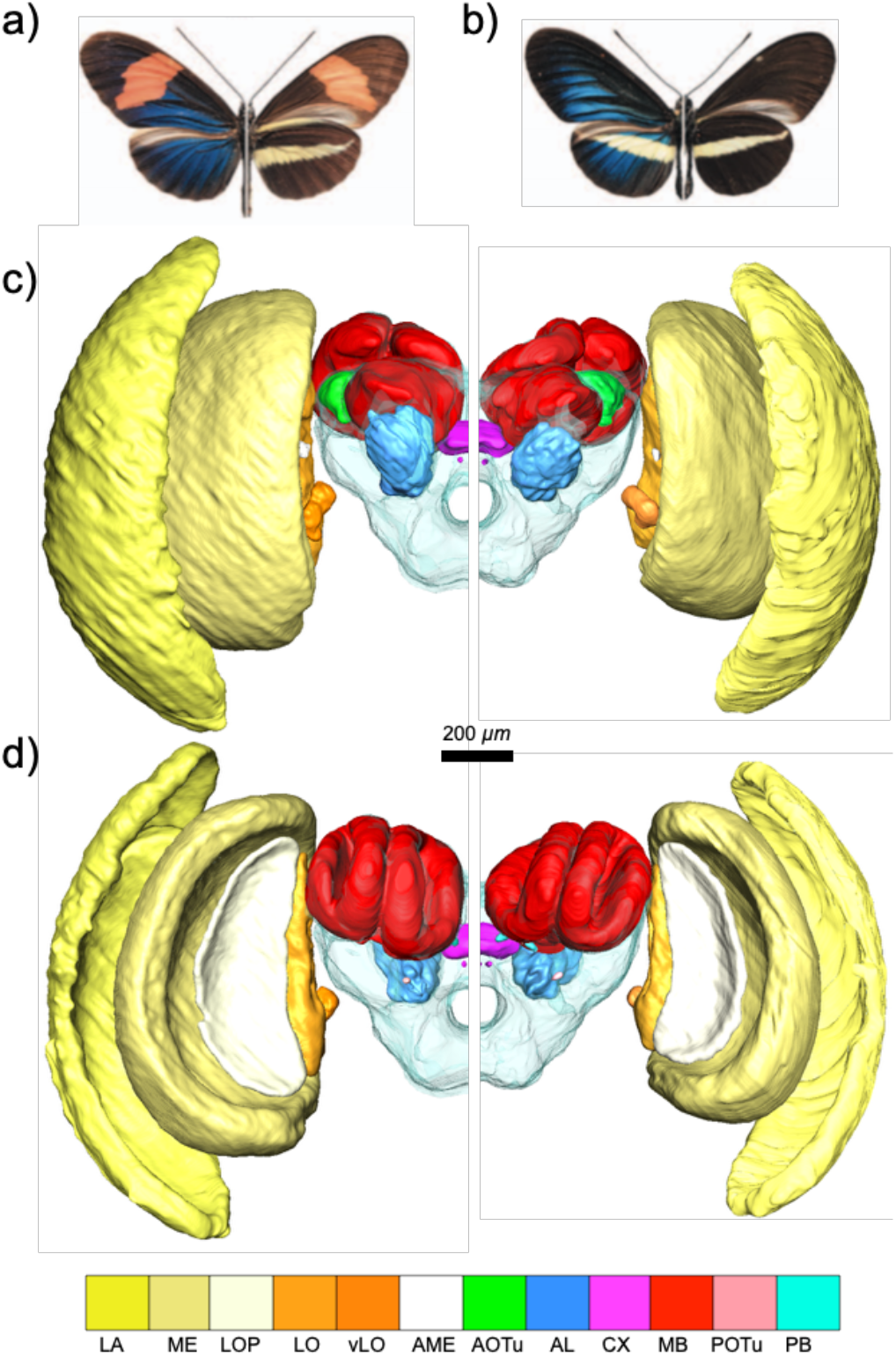
Upper row. Dorsal (left side) and ventral (right-side) overview of the wing colour patterns of a) *H. e. venus* and b) *H. chestertonii*. Middle and bottom rows. Rostral (c) and caudal (d) view of volumetric reconstructions of the brains of wild male *H. e. venus* (left half) and *H. chestertonii* (right half). The brain images showcase the rCBR (transparent), the lamina (LA), the medulla (ME), the lobula plate (LOP), the lobula (LO), the ventral lobula (vLO), the accessory medulla (AME), the anterior optic tubercle (AOTu), the posterior optic tubercle (POTu), the mushroom bodies (MB), the protocerebral bridge (PB), the central complex (CX) and the antennal lobe (AL).

### iv) Statistical analyses

We used linear mixed effects models (lmer function in the *lmer* package in R (Bates et al., 2014)) to assess the scaling relationships between the neuropils of interest and an independent measure of brain size. We designated rCBR as the independent allometric control after determining that it does not vary significantly between sexes (n = 36, F= 0.7664; p=0.387) or species (n = 36, F= 0.0006; p=0.980). In our models, the volume of each neuropil was a function of the fixed factors “rCBR” and “species”, as well as the random factor “sex”. The effect of “species” was determined by comparing model performance with and without the term via Wald 𝒳^2^ tests (*Anova* function in the R package *CAR* (Fox & Weisberg, 2018)).

We next implemented standardised major axis regression analyses (SMATR) to confirm non-allometric shifts in brain structure across wild caught *H. e. venus* and *H. chestertonii* (*SMATR* package in R (Warton et al., 2012)). These analyses test for significant between-groups shifts in the allometric relationship log *y*=*β*log *x* + *α*, which describes the association between a given trait (*y*, i.e. volume of neuropil of interest) and an independent measure of size or allometric control (*x*, i.e. volume of rCBR). *β* is the slope and *α* is the elevation of the line of fit. Shifts in slope reflect different scaling relationships between the species being compared, whereas changes in the intercept or elevation (grade-shifts) are interpreted as non-allometric divergence (Kruska, 2005), both of which may result from mosaic brain evolution (Barton & Harvey, 2000). Major axis-shifts on the other hand occur along a shared allometry and underly changes in neuropil volume in concert with the rest of the brain (Barton & Harvey, 2000; Montgomery, Mundy, et al., 2016). We additionally performed these tests separately in insectary individuals reared in common garden conditions to determine the extent to which between-species differences result from environmental effects on brain structure.

We created a combined dataset with volumetric brain measurements of *H. e. cyrbia, H. himera, H. e. venus* and *H. chestertonii* to test for shared patterns of ecologically driven divergence. The habitat of each species has been previously designated as low- (*H. e. cyrbia* and *H. e. venus*) or high-altitude (*H. himera* and *H. chestertonii*) based on patters of macroclimatic variation across their distribution (e.g. high-altitude habitats show characteristically lower annual mean temperatures, lower rainfall and greater precipitation seasonality) (Rivas-Sánchez, Melo-Flórez, et al., 2023). We used linear mixed models where variation in neuropil volume was partitioned in that attributable to the fixed factors “habitat type” (low or high-altitude), “locality” (Colombia or Ecuador) and their interaction. Comparing models with and without these terms allowed detecting significant effects on the volume of specific neuropils stemming from shared sources of ecologically based selection (“habitat type”), unique histories of each divergence event (“locality”) and unique evolutionary responses to shared habitat shifts (“interaction”) (Langerhans & DeWitt, 2004). These models included our allometric control (“rCBR”) as a fixed factor and “sex” as random term. The relative contribution of each factor to trait variation was subsequently estimated using semi-partial R^2^ measures using the *r2Beta* function (Kenward Roger method in the R package *r2glmm* (Jaeger et al., 2017)). Brain data, detailed test results and annotated R scripts used here are provided as supplementary material (supplementary tables S0-S1).

## Results

### i) Heritable divergence in sensory brain structures

Likelihood-rate model comparisons revealed differences between wild *H. e. venus* and *H. chestertonii* in the volume of the medulla (n = 36, 𝒳1^2^= 5.802; p<0.05, the lobula (n = 36, 𝒳1^2^= 9.7488; p<0.01), the lobula plate (n = 36, 𝒳1^2^= 5.592; p<0.05) and the antennal lobe (n = 36, 𝒳1^2^= 6.1646; p<0.01), which are generally larger in *H. e. venus* compared to *H. chestertonii*. SMATR analyses confirmed that this variation is driven by non-allometric grade-shifts (Figure 2A-D, supplementary table S2) affecting specific optic lobe neuropils (medulla, Wald 𝒳1^2^ = 5.262, *P* < 0.05; lobula, Wald 𝒳1^2^ = 7.004, *P* < 0.01 and lobula plate, Wald *χ*^2^1 = 4.553, *P* < 0.05) and the antennal lobe (Wald *χ*^2^1 = 5.371, *P* < 0.05).

**Figure 2.**
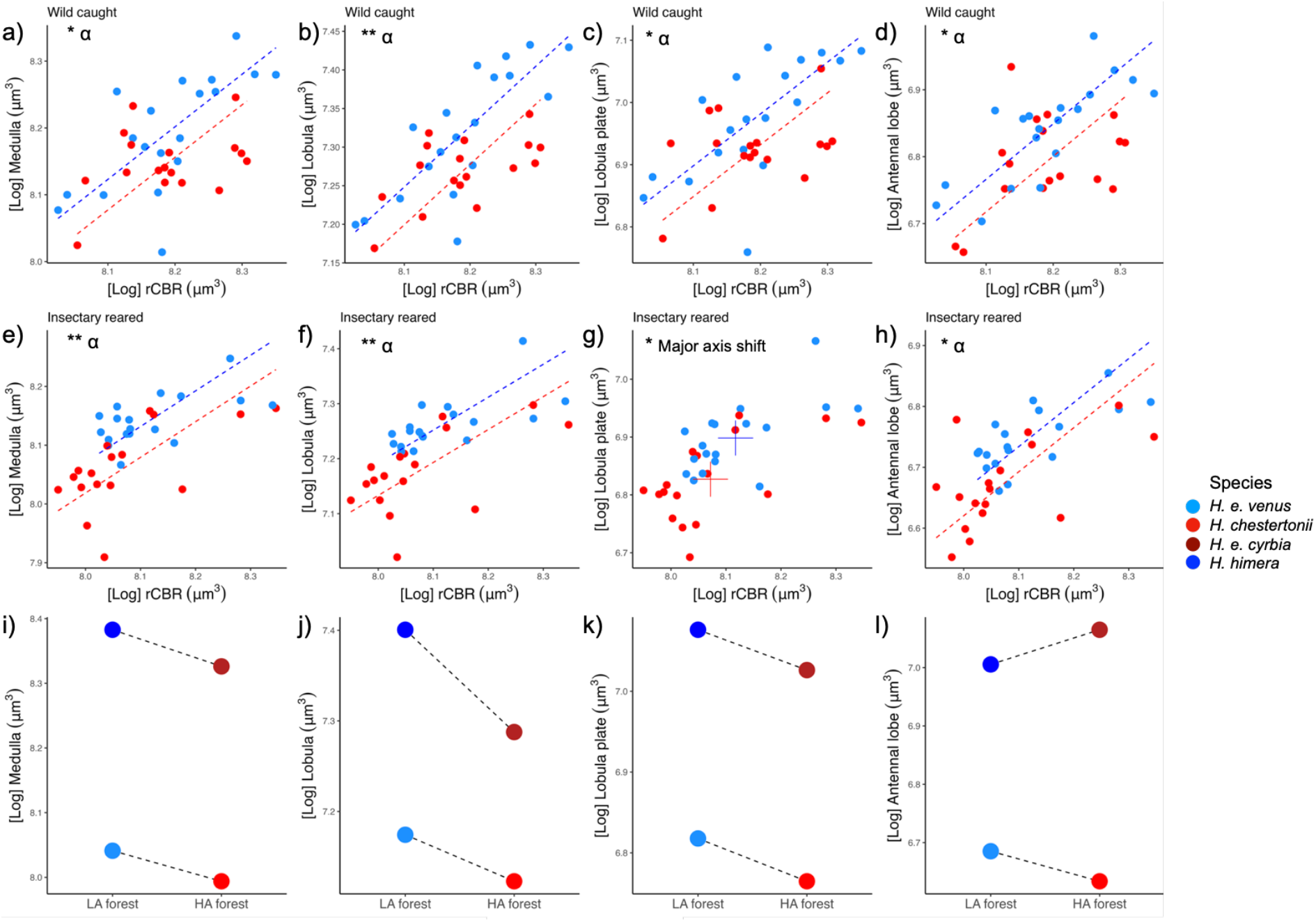
First row. Grade (α)-shifts between wild *H. chestertonii* and *H. e venus* in scaling relationships against the rCBR of the a) medulla, b) lobula, c) lobula plate and d) antennal lobe). Second row. Grade (α)-shifts (e, f and d) and major-axis (g) shift between insectary-reared *H. chestertonii* and *H. e venus* in the scaling relationships against the rCBR of the e) medulla, f) lobula, c) lobula plate and d) antennal lobe). For all the plots, points and dashed lines represent, respectively, individuals and allometries between neuropil and rCBR volume for *H. chestertonii* (red) and *H. e. venus* (blue). For plot g, + symbols represent the centroid of *H. chestertonii* (red) and *H. e. venus* (blue). Asterisks convey significance levels of Grade (α) and major axis shifts (likelihood-ratio statistic). * p < 0.05. ** p < 0.01. Third row. Levels of parallelism in *Heliconius* species pairs in the relative volume of the i) medulla, j) lobula, k) lobula plate and l) antennal lobe. Coloured dots convey the volume of neuropils for butterflies with a common rCBR size. Dashed lines connect species from the same locality and different forest types. In all the plots, *H. chestertonii* is in red, *H. e. venus* in blue, *H. himera* in firebrick and *H. e. cyrbia* in dodger blue. LA= low-altitude forest and HA=high-altitude forest.

This pattern was preserved in our sample of insectary-reared individuals (grade-shifts in the medulla, Wald *χ*^2^1 = 7.629, *P* < 0.01; lobula, Wald 𝒳1^2^ = 9.046, *P* < 0.01 and antennal lobe Wald *χ*^2^1 = 4.42, *P* < 0.05), with the exception of the lobula plate, that instead exhibited a major-axis shift between species (Wald 𝒳1^2^ = 6.999, *P* < 0.01), along with several other visual structures (ventral lobula, Wald 𝒳1^2^ = 5.956, *P* < 0.05; lamina, Wald 𝒳1^2^ = 6.123, *P* < 0.05 and AOTu, Wald 𝒳1^2^ = 6.611, *P* < 0.05, Figure 2E-H, supplementary Table S3). Further comparisons showed that variation between insectary-reared and wild *H. chestertonii* are mostly explained by allometric scaling, with no significant *β* or *α* shifts except for the POTu (Wald 𝒳1^2^ = 23.05, *P* < 0.01, Supplementary Table S4), whereas insectary-reared *H. e. venus* showed non-allometric reductions in the size of the medulla, the lobula and the antennal lobe compared to wild conspecifics (Supplementary Table S5).

### ii) Consistent shifts in visual investment during parallel speciation

For the brain regions identified as divergent in wild-caught individuals, we compared linear mixed models explaining variation in neuropil size across *H. e. venus, H. chestertonii, H. e. cyrbia*, and *H. himera*. We found significant “habitat type” effects on the medulla (𝒳1^2^ = 20.849, *P* < 0.001, semi-partial R^2^= 0.246), the lobula (𝒳1^2^ = 30.905, *P* < 0.001, R^2^= 0.494) and the lobula plate (𝒳1^2^ = 13.911, *P* < 0.001, R^2^= 0.174), which, for a given rCBR size, are significantly smaller in the high-altitude specialists *H. chestertonii* and *H. himera* than in their parapatric lowland *H. erato* populations (Figure 2I-K). We also identified significant “locality” effects on the medulla (𝒳1^2^ = 426.233, *P* < 0.001), the lobula (𝒳1^2^= 135.831, *P* < 0.001), the lobula plate (𝒳1^2^ = 173.746, *P* < 0.001) and the antennal lobe (𝒳1^2^ = 386.4651, *P* < 0.001), all relatively larger in *H. e. cyrbia* and *H. himera* compared to the Colombian populations. In addition, we found a significant effect of the habitat type x locality interaction on the antennal lobe (𝒳1^2^ = 15.4913, *P* < 0.001), consistent with contrasting effects of high-altitude in its volume across Colombia and Ecuador (Figure 2L).

## Discussion

We leveraged early speciation events *Heliconius* species to test for adaptive shifts in neuroanatomy associated with recent ecological divergence across forest types. First, we found that wild *H. chestertonii* individuals sampled from their high-altitude forest habitats in Colombia show non-allometric reductions in the size of specific optic lobe neuropils compared to *H. e. venus* butterflies captured in lower altitude forests. Next, by rearing these same species under common garden conditions, we determined that heritable variation largely underlies this pattern of divergence. In addition, the results of our analyses incorporating additional data on wild *H. e. cyrbia* and *H. himera* demonstrate that optic lobe size variation is at least partially attributable to shared selection regimes across the geographically separated high-altitude forests inhabited by *H. himera* and *H. chestertonii*, likely related to parallel adaptation in similar visual environments. In contrast, the antennal lobe shows disparate trait-shifts across *H. himera* and *H. chestertonii*, suggesting different responses to ostensibly equivalent habitat-shifts and independence between brain regions associated with separate sensory modalities.

Conserved scaling relationships between brain components are thought to reflect the effects of stabilizing selection limiting the independent evolution of functionally linked brain regions (Montgomery, Mundy, et al., 2016). Mosaic-like patterns of divergence in brain structure, on the other hand, suggest selection targeting specific brain regions producing adaptive behavioural shifts (Barton & Harvey, 2000). Via model comparisons, we detected significant levels of between-species variation in the size of multiple sensory neuropils in *H. e. venus* and *H. chestertonii*. SMATR regressions further showed that these differences largely consist of grade-shifts that affect the medulla, the lobula, the lobula plate and the antennal lobe, resulting in those neuropils being smaller in wild *H. chestertonii* relative to *H. e. venus* for a given brain size. These grade-shifts between species were also detected in the medulla, the lobula and the antennal lobe of butterflies reared in a common garden, where altitude and climatic conditions were similar to those naturally experienced by *H. e. chestertonii* (Rivas-Sánchez, Melo-Flórez, et al., 2023). The lobula plate, instead, showed a major axis-shift that nevertheless, results in this neuropil being functionally smaller in insectary reared *H. e. chestertonii*. Thus, it is likely that to a large extent, the patterns of brain divergence we observed in wild samples represent heritable variation rather than environmentally induced responses. Importantly, insectary-reared *H. chestertonii* exhibited major axis-shifts in these neuropils compared to wild-caught conspecifics that can be explained by concerted reductions in overall brain size, expected from contrasting foraging experiences between forests and insectary cages (Montgomery, Merrill, et al., 2016). However, although an equivalent comparison in *H. e. venus* revealed similar overall brain size reductions in insectary-reared butterflies, this group also showed non-allometric shifts relative to wild conspecifics that bring it closer to a *H. chestertonii*-like phenotype (e.g. grade-shifts reducing the volume of the antennal lobe). A comparison between *H. e. venus* siblings exposed to high and low-altitude habitats would help determine the potential of brain plasticity as “pacemaker” mechanism facilitating non-allometric shifts in these sensory neuropils (Levis & Pfennig, 2016).

In the independent high-altitude specialist *H. himera*, multiple optic lobe neuropils are reduced compared to the same structures in *H. e. cyrbia* from lower altitude (Montgomery & Merrill, 2017). Our results show a parallel pattern of variation in three optic lobe neuropils in *H. e. venus* and *H. chestertonii*, which are segregated in ecologically similar forest types at low and high-altitude respectively (Rivas-Sánchez, Melo-Flórez, et al., 2023). These include the medulla, which in *Drosophila* and honeybees has been implicated in light parallelization, colour vision and motion detection (Borst, 2009; Morante & Desplan, 2004; Paulk et al., 2009), as well as the lobula and the lobula plate, which are involved in escape/chase reactions to visual stimuli and motion computation (Hausen, 1984). Our model comparisons incorporating data from the four species favour divergent selection as a factor explaining these repeated trends of optic lobe size reduction across the Ecuador and Colombian localities. In particular, “habitat type” explains 17.4-49.4 % of the overall neuropil size variation in these species, implying independent, repeated trait-shifts that are consistent with adaptation shaped by similar ecological conditions (Bolnick et al., 2018; Langerhans & DeWitt, 2004). This reflects patterns seen in the more distantly related species pair *H. cydno*-*H. melpomene*, where habitat partitioning along gradients of increasing light intensity in forest interior-forest edge axes is accompanied by non-neutral reductions in the size of the optic lobe and the anterior optic tubercle, as well as lower number of eye facets and reduced corneal area (Montgomery et al., 2021; Seymoure et al., 2015; Wright et al., 2023).

Perceiving and processing visual information is energetically expensive (Laughlin et al., 1998) and these particularly high metabolic demands are met with selection for efficiency, ultimately regulating the design of sensory and neural systems (Laughlin, 2001; Laughlin et al., 1998; Niven & Laughlin, 2008). For instance, in *Drosophila* populations reared in captivity, selection for finding resources using visual cues is relaxed relative to natural conditions, while selective pressures for low energy consumption are maintained, leading to the evolution of smaller (*i*.*e*. less energetically demanding) eye size over few generations (Tan et al., 2005). Low-altitude forests in Colombia and Ecuador are relatively dim due to their dense vegetation, forest structure and lower levels of solar radiation, (Dell’Aglio et al., 2022; Rivas-Sánchez, Gantiva-Q, et al., 2023; Rivas-Sánchez, Melo-Flórez, et al., 2023), which likely results in darker and more visually complex light-environments compared to high-altitude forests. Thus, the shifts towards a reduced volume in some optic lobe neuropils in *H. chestertonii* and *H. himera* could potentially be consequence of sustained selection for low energetic costs matched with more lenient vision requirements in their high-altitude habitats. These putative local adaptations may in turn hinder gene flow in the absence of geographic barriers, via lower rates of heterospecific encounters between resident individuals and migrants, or hybrids with fitness deficits (Coyne & Orr, 2004; Montgomery et al., 2021; Nosil et al., 2005).

In contrast to consistent changes in the visual neuropils, we identified incongruent evolutionary responses in the antennal lobe between Colombian and Ecuadorian localities. Among Lepidoptera, variation in the relative size of the antennal lobe is assumed to reflect different emphasis on olfactory cues between species in habitats with contrasting availability of visual information (Montgomery & Ott, 2015; Morris et al., 2021; Stöckl et al., 2016). While investment in this neuropil is higher in *H. himera* in respect to *H. e. cyrbia* (Montgomery & Merrill, 2017), here, we show it is lower in *H. chestertonii* compared to its lowland counterpart *H. e. venus*, entailing locality specific responses to habitat shifts towards high-altitude and independent evolution of brain components linked to separate vision and olfaction without apparent developmental trade-offs between investment on either of these sensory modalities (Farnworth & Montgomery, 2022). Variation in the antennal lobe is heritable in *H. e. venus* and *H. chestertonii*, but not in *H. e. cyrbia* and *H. himera*, therefore, a possibility is that the evolution of this neuropil is influenced by lasting effects of recent geneflow with other *Heliconius* species or genetic drift (Moore & Hendry, 2005; Oke et al., 2017; Stuart et al., 2017). Alternatively, the habitat-categorizations employed here might not capture aspects of environmental variation that may exert locality specific selective pressures (Bolnick et al., 2018; Stuart et al., 2017) across the habitats of *H. himera* and *H. chestertonii*, resulting in disparate trait-shifts across these species. For instance, relative humidity and rainfall are significantly lower in the habitat of *H. himera* compared to the forests inhabited by *H. chestertonii* (Rivas-Sánchez, Melo-Flórez, et al., 2023). Further research could explore whether habitat differences such as these affect the sensory background under which olfactory cues are perceived, which may in turn relate to the patterns of variation in the antennal lobe we observed across our focal, independent high-altitude *Heliconius* species (Dell’Aglio et al., 2022; Montgomery & Merrill, 2017)

In summary, our study presents evidence of heritable neuroanatomical divergence between two closely related species from the *H. erato* complex that are segregated in forests at different altitude in Colombia. Similar to physiological, life-history and behavioural traits (Rivas-Sánchez, Gantiva-Q, et al., 2023; Rivas-Sánchez, Melo-Flórez, et al., 2023), some of these brain-shifts are mirrored in a separate pair of species that occupy similar ecological niches in Ecuador, suggesting local adaptations shaped by shared sensory environments. Habitat variation between different forest types likely results in fitness deficits in migrants with alternative brain phenotypes, facilitating species divergence in the face of ongoing geneflow. However, these evolutionary responses are not perfectly parallel, suggesting lineage specific evolutionary responses driven by cryptic sources of environmental variation.

## Author contributions

SHM and RMM conceptualised the study. DFRS performed the data collection under the supervision and support of CPD, CS, RMM and SHMM. DFRS curated the data. SHM, RMM and DFRS designed the analyses. DFRS performed the analyses and wrote a first draft, which was subsequently edited by all the authors.

## Acknowledgements

We are thankful to Benito Wainwright, Amaia Alcalde and Max Farnworth from the EBaB lab for valuable suggestions and comments on the analyses performed here. Additional thanks to Isabel León, Lina Melo and Andrea Aragón for assistance with growing *Passiflora* plants and rearing insectary butterflies as well as Nicol Rueda and Lucas Barrientos for help during field collections. Rearing and field collections were conducted under the permit no. 530 issued by the Autoridad Nacional de Licencias Ambientales of Colombia (ANLA). This study was funded by a Natural Environment Research Council Independent Research Fellowship (NE/N014936/1) to SHM, a European Research Council Starter Grant (851040) to RMM, and a scholarship to DFRS from the Ministerio de Ciencia Tecnología e Innovación de Colombia (Convocatoria No. 860).

## Data availability

Supplementary information and raw data can be found in Supplementary Tables S0– S5 and annotated R scripts. Files are available in the Dryad Digital Repository.

