## Supplementary material for "Repeated evolution of reduced visual investment at the onset of ecological speciation in high-altitude *Heliconius* butterflies"

---
output:
  pdf_document: default
  html_document: default
---

```
#Importing and sorting the data

WildBrains<-subset(AllBrains, Origin=="wild")
WildFemales<-subset(AllBrains, Origin=="wild" & Sex=="female")
WildMales<-subset(AllBrains, Origin=="wild" & Sex=="male")
InsectaryBrains<-subset(AllBrains, Origin=="insectary")
InsectaryFemales<-subset(AllBrains, Origin=="insectary" & Sex=="female")
InsectaryMales<-subset(AllBrains, Origin=="insectary" & Sex=="male")
FemaleBrains<-subset(AllBrains, Sex=="female")
MaleBrains<-subset(AllBrains, Sex=="male")
Chestertoniibrains<-subset(AllBrains, Subspecies=="chestertonii")
VenusBrains<-subset(AllBrains, Subspecies=="venus")
Wildchestertonii<-subset(AllBrains, Subspecies=="chestertonii" & Origin=="wild")
Wildvenus<-subset(AllBrains, Subspecies=="venus"& Origin=="wild")
Insectarychestertonii<-subset(AllBrains, Subspecies=="chestertonii" &
Origin=="insectary")
Insectaryvenus<-subset(AllBrains, Subspecies=="venus"& Origin=="insectary")

summary(AllBrains)

#Checking for sex and subspecies effects on the allometric control (there are
neither)

RCBRreducedw<-lm(RCBR_log ~Subspecies+Sex, data = WildBrains)
RCBRaltw<-lm(RCBR_log ~Subspecies, data = WildBrains)
RCBRaltw1<-lm(RCBR_log ~Sex, data = WildBrains)

Anova(RCBRreducedw)
anova(RCBRreducedw,RCBRaltw1)
anova(RCBRreducedw,RCBRaltw)


summary(RCBRreducedw)
summary(RCBRaltw)
summary(RCBRaltw1)


#Medulla Modelling (Subspecies*)


MedullaFull<-lmer(ME_log ~ RCBR_log *Subspecies+(1|Sex), data = WildBrains)
summary(MedullaFull)
```

```
drop1(MedullaFull, test="Chisq")
newmedulla<-update(MedullaFull, ~. -RCBR_log:Subspecies)

drop1(newmedulla, test="Chisq")

MedullaReduced<-lmer(ME_log ~ RCBR_log +Subspecies+(1|Sex), data =
WildBrains)
MedullaAlt<-lmer(ME_log ~ RCBR_log+(1|Sex), data = WildBrains)

anova(MedullaReduced, MedullaAlt)
Anova(MedullaReduced)

summary(MedullaReduced)

#Lobulla Modelling (Subspecies*)


LobulaFull<-lmer(LO_log ~ RCBR_log *Subspecies +(1|Sex), data = WildBrains)
summary(LobulaFull)

drop1(LobulaFull, test="Chisq")
newlobula<-update(LobulaFull, ~. -RCBR_log:Subspecies)

drop1(newlobula, test="Chisq")

LobulaReduced<-lmer(LO_log ~ RCBR_log +Subspecies +(1|Sex), data =
WildBrains)
LobulaAlt<-lmer(LO_log ~ RCBR_log+(1|Sex), data = WildBrains)

anova(LobulaReduced, LobulaAlt)
Anova(LobulaReduced)


summary(LobulaReduced)



#Lobulla plate models (Subspecies*)

LobulapFull<-lmer(LOP_log ~ RCBR_log *Subspecies +(1|Sex), data = WildBrains)
summary(LobulapFull)


drop1(LobulapFull, test="Chisq")
newlobulap<-update(LobulaFull, ~. -RCBR_log:Subspecies)

drop1(newlobulap, test="Chisq")
```

```
LobulaPReduced<-lmer(LOP_log ~ RCBR_log +Subspecies+(1|Sex), data =
WildBrains)
LobulapAlt<-lmer(LOP_log ~ RCBR_log+(1|Sex), data = WildBrains)

anova(LobulaPReduced, LobulapAlt)
Anova(LobulaPReduced)

summary(LobulaPReduced)

#Ventral lobula modelling ()


vLobulaFull<-lmer(vLO_log ~ RCBR_log *Subspecies +(1|Sex), data = WildBrains)
summary(vLobulaFull)

drop1(vLobulaFull, test="Chisq")
newvlobula<-update(vLobulaFull, ~. -RCBR_log:Subspecies)

drop1(newvlobula, test="Chisq")
newvlobula1<-update(newvlobula, ~. -Subspecies)

drop1(newvlobula1, test="Chisq")


vLobulaReduced<-lmer(vLO_log ~ RCBR_log+(1|Sex), data = WildBrains)
vLobulaAlt<-lmer(vLO_log ~ RCBR_log+Subspecies+(1|Sex), data = WildBrains)

anova(vLobulaReduced, vLobulaAlt)
Anova(vLobulaAlt)

summary(vLobulaReduced)




#Lamina modelling ()

LaminaFull<-lmer(LA_log ~ RCBR_log *Subspecies +(1|Sex), data = WildBrains)
summary(LaminaFull)


drop1(LaminaFull, test="Chisq")
newlamina<-update(LaminaFull, ~. -RCBR_log:Subspecies)

drop1(newlamina, test="Chisq")
newlamina1<-update(newlamina, ~. -Subspecies)

drop1(newlamina1, test="Chisq")
```

```
LaminaReduced<-lmer(LA_log ~ RCBR_log+Subspecies+(1|Sex), data =
WildBrains)
LaminaAlt<-lmer(LA_log ~ RCBR_log +(1|Sex), data = WildBrains)

anova(LaminaReduced, LaminaAlt)
Anova(LaminaReduced)


summary(LaminaReduced)




#Accessory medulla modelling ()

ameFull<-lmer(AME_log ~ RCBR_log *Subspecies +(1|Sex), data = WildBrains)
summary(ameFull)

drop1(ameFull, test="Chisq")
newame<-update(ameFull, ~. -RCBR_log:Subspecies)

drop1(newame, test="Chisq")
newame1<-update(newame, ~. -Subspecies)

drop1(newame1, test="Chisq")


ameReduced<-lmer(AME_log ~ RCBR_log+(1|Sex), data = WildBrains)
ameAlt<-lmer(AME_log ~ RCBR_log+Subspecies+(1|Sex), data = WildBrains)

anova(ameReduced, ameAlt)
Anova(ameAlt)

summary(ameReduced)


#Posterior optic tubercle modelling (no allometric growth)

potuFull<-lmer(POTu_log ~ RCBR_log *Subspecies +(1|Sex), data = WildBrains)
summary(potuFull)

drop1(potuFull, test="Chisq")
newpotu<-update(potuFull, ~. -RCBR_log:Subspecies)

drop1(newpotu, test="Chisq")
newpotu1<-update(newpotu, ~. -Subspecies)

drop1(newpotu1, test="Chisq")
```

```
potuReduced<-lmer(POTu_log ~ RCBR_log+(1|Sex), data = WildBrains)
potuAlt<-lmer(POTu_log ~ RCBR_log+Subspecies+(1|Sex), data = WildBrains)

anova(potuReduced, potuAlt)
Anova(potuAlt)

summary(potuReduced)

#Anterior optic tubercle modelling ()

AOTuFull<-lmer(AOTu_log ~ RCBR_log *Subspecies +(1|Sex), data = WildBrains)
summary(AOTuReduced)


drop1(AOTuFull, test="Chisq")
newAOTu<-update(AOTuFull, ~. -RCBR_log:Subspecies)

drop1(newAOTu, test="Chisq")
newAOTu1<-update(newAOTu, ~. -Subspecies)

drop1(newAOTu1, test="Chisq")


AOTuReduced<-lmer(AOTu_log ~ RCBR_log+(1|Sex), data = WildBrains)
AOTuAlt<-lmer(AOTu_log ~ RCBR_log+Subspecies+(1|Sex), data = WildBrains)

anova(AOTuReduced, AOTuAlt)

summary(AOTuAlt)


#Antennal Lobe models (Subspecies*)

alobeFull<-lmer(AL_log ~ RCBR_log *Subspecies +(1|Sex), data = WildBrains)
summary(alobeFull)


drop1(alobeFull, test="Chisq")
alobenew<-update(alobeFull, ~. -RCBR_log:Subspecies)

drop1(alobenew, test="Chisq")


alobeReduced<-lmer(AL_log ~ Subspecies+RCBR_log+(1|Sex), data = WildBrains)
alobeAlt<-lmer(AL_log ~ RCBR_log+(1|Sex), data = WildBrains)

anova(alobeReduced, alobeAlt)
Anova(alobeReduced)
```

```
summary(alobeReduced)



#Exploratory plots to visually inspect scaling relationships

#Wild caught animals grouped as subspecies


medullawildplot<-ggplot(WildBrains, aes(x=RCBR_log, y=ME_log,
color=Subspecies)) + geom_point()+labs(x = bquote('[Log] rCBR'~(µm^3)), y =
bquote('[Log] Medulla'~(µm^3)))  + theme(plot.title = element_text(hjust = 0.5))
+theme_classic()+geom_point(size=3)+ scale_color_manual(values = c("red",
"dodgerblue"))+theme(axis.title = element_text(size = 15))+ggtitle("Wild caught")
lobulawildplot<-ggplot(WildBrains, aes(x=RCBR_log, y=LO_log, color=Subspecies))
+ geom_point()+labs(x = bquote('[Log] rCBR'~(µm^3)), y = bquote('[Log]
Lobula'~(µm^3))) + theme(plot.title = element_text(hjust = 0.5))+theme_classic()
+geom_point(size=3)+ scale_color_manual(values = c("red", "dodgerblue"))
+theme(axis.title = element_text(size = 15))+ggtitle("Wild caught")
lobulaplatewildplot<-ggplot(WildBrains, aes(x=RCBR_log, y=LOP_log,
color=Subspecies)) + geom_point()+labs(x = bquote('[Log] rCBR'~(µm^3)), y =
bquote('[Log] Lobula plate'~(µm^3)))  + theme(plot.title = element_text(hjust = 0.5))
+theme_classic()+geom_point(size=3)+ scale_color_manual(values = c("red",
"dodgerblue"))+theme(axis.title = element_text(size = 15))+ggtitle("Wild caught")
vlobulawildplot<-ggplot(WildBrains, aes(x=RCBR_log, y=vLO_log,
color=Subspecies)) + geom_point()+labs(x = bquote('[Log] rCBR'~(µm^3)), y =
bquote('[Log] Ventral lobula'~(µm^3))) + theme(plot.title = element_text(hjust = 0.5))
+theme_classic()+geom_point(size=3)+ scale_color_manual(values = c("red",
"dodgerblue"))+theme(axis.title = element_text(size = 15))
amedullawildplot<-ggplot(WildBrains, aes(x=RCBR_log, y=AME_log,
color=Subspecies)) + geom_point()+labs(x = bquote('[Log] rCBR'~(µm^3)), y =
bquote('[Log] Accessory medulla'~(µm^3))) + ggtitle("Wild caught") + theme(plot.title
= element_text(hjust = 0.5))+theme_classic()+geom_point(size=3)+
scale_color_manual(values = c("red", "dodgerblue"))+theme(axis.title =
element_text(size = 15))
aotuberclewildplot<-ggplot(WildBrains, aes(x=RCBR_log, y=AOTu_log,
color=Subspecies)) + geom_point()+labs(x = bquote('[Log] rCBR'~(µm^3)), y =
bquote('[Log] Anterior optic tubercle'~(µm^3))) + ggtitle("Wild caught") +
theme(plot.title = element_text(hjust = 0.5))+theme_classic()+geom_point(size=3)+
scale_color_manual(values = c("red", "dodgerblue"))+theme(axis.title =
element_text(size = 15))
alobewildplot<-ggplot(WildBrains, aes(x=RCBR_log, y=AL_log, color=Subspecies))
+ geom_point()+labs(x = bquote('[Log] rCBR'~(µm^3)), y = bquote('[Log] Antennal
lobe'~(µm^3))) + theme(plot.title = element_text(hjust = 0.5))+theme_classic()
+geom_point(size=3)+ scale_color_manual(values = c("red", "dodgerblue"))
+theme(axis.title = element_text(size = 15))+ggtitle("Wild caught")
laminawildplot<-ggplot(WildBrains, aes(x=RCBR_log, y=LA_log, color=Subspecies))
+ geom_point()+labs(x = bquote('[Log] rCBR'~(µm^3)), y = bquote('[Log]
Lamina'~(µm^3))) + ggtitle("Wild caught") + theme(plot.title = element_text(hjust =
```

```
0.5))+theme_classic()+geom_point(size=3)+ scale_color_manual(values = c("red", "dodgerblue"))+theme(axis.title = element_text(size = 15))
potuberclewildplot<-ggplot(WildBrains, aes(x=RCBR_log, y=POTu_log, color=Subspecies)) + geom_point()+labs(x = bquote('[Log] RoCB'~(µm^3)), y = bquote('[Log] Posterior optic tubercle'~(µm^3))) + ggtitle("Wild caught") + theme(plot.title = element_text(hjust = 0.5))+theme_classic()+geom_point(size=3)+ scale_color_manual(values = c("red", "dodgerblue"))+theme(axis.title = element_text(size = 15))

#Wild males grouped as species

medullawildmaleplot<-ggplot(WildMales, aes(x=RCBR_log, y=ME_log, color=Subspecies)) + geom_point()+labs(y= "Medulla [Log]", x = "Rest of the Central Brain [Log]") + ggtitle("Wild males") + theme(plot.title = element_text(hjust = 0.5))+theme_classic()+geom_point(size=3)+ scale_color_manual(values = c("firebrick", "dodgerblue"))
ggplot(WildMales, aes(x=RCBR_log, y=LO_log, color=Subspecies)) + geom_point()+labs(y= "Lobula [Log]", x = "Rest of the Central Brain [Log]") + ggtitle("Wild males") + theme(plot.title = element_text(hjust = 0.5))+theme_classic()+geom_point(size=3)+ scale_color_manual(values = c("firebrick", "dodgerblue"))
ggplot(WildMales, aes(x=RCBR_log, y=LOP_log, color=Subspecies)) + geom_point()+labs(y= "Lobula plate [Log]", x = "Central Brain (minus AL & AOTu) [Log]") + ggtitle("Wild males") + theme(plot.title = element_text(hjust = 0.5))+theme_classic()+geom_point(size=3)+ scale_color_manual(values = c("firebrick", "dodgerblue"))
ggplot(WildMales, aes(x=RCBR_log, y=vLO_log, color=Subspecies)) + geom_point()+labs(y= "Ventral lobulla [Log]", x = "Central Brain (minus AL & AOTu) [Log]") + ggtitle("Wild males") + theme(plot.title = element_text(hjust = 0.5))+theme_classic()+geom_point(size=3)+ scale_color_manual(values = c("firebrick", "dodgerblue"))
ggplot(WildMales, aes(x=RCBR_log, y=AME_log, color=Subspecies)) + geom_point()+labs(y= "Accessory medulla [Log]", x = "Central Brain (minus AL & AOTu) [Log]") + ggtitle("Wild males") + theme(plot.title = element_text(hjust = 0.5))+theme_classic()+geom_point(size=3)+ scale_color_manual(values = c("firebrick", "dodgerblue"))
ggplot(WildMales, aes(x=RCBR_log, y=AOTu_log, color=Subspecies)) + geom_point()+labs(y= "Anterior optic tubercle [Log]", x = "Central Brain (minus AL & AOTu) [Log]") + ggtitle("Wild males") + theme(plot.title = element_text(hjust = 0.5))+theme_classic()+geom_point(size=3)+ scale_color_manual(values = c("firebrick", "dodgerblue"))
ggplot(WildMales, aes(x=RCBR_log, y=AL_log, color=Subspecies)) + geom_point()+labs(y= "Antennal lobe [Log]", x = "Central Brain (minus AL & AOTu) [Log]") + ggtitle("Wild males") + theme(plot.title = element_text(hjust = 0.5))+theme_classic()+geom_point(size=3)+ scale_color_manual(values = c("firebrick", "dodgerblue"))
ggplot(WildMales, aes(x=RCBR_log, y=LA_log, color=Subspecies)) + geom_point()+labs(y= "Lamina [Log]", x = "Central Brain (minus AL & AOTu) [Log]") + ggtitle("Wild males") + theme(plot.title = element_text(hjust = 0.5))+theme_classic()+geom_point(size=3)+ scale_color_manual(values = c("firebrick", "dodgerblue"))

#Wild females separated by species
```

```
ggplot(WildFemales, aes(x=RCBR_log, y=ME_log, color=Subspecies)) +
geom_point()+labs(y= "Medulla [Log]", x = "Central Brain (minus AL & AOTu) [Log]")
+ ggtitle("Wild females") + theme(plot.title = element_text(hjust = 0.5))
+theme_classic()+geom_point(size=3)+ scale_color_manual(values = c("firebrick",
"dodgerblue"))
ggplot(WildFemales, aes(x=RCBR_log, y=LO_log, color=Subspecies)) +
geom_point()+labs(y= "Lobula [Log]", x = "Central Brain (minus AL & AOTu) [Log]") +
ggtitle("Wild females") + theme(plot.title = element_text(hjust = 0.5))+theme_classic()
+geom_point(size=3)+ scale_color_manual(values = c("firebrick", "dodgerblue"))
lobulaplatewildfemaleplot<-ggplot(WildFemales, aes(x=RCBR_log, y=LOP_log,
color=Subspecies)) + geom_point()+labs(y= "Lobula plate [Log]", x = "Central Brain
(minus AL & AOTu) [Log]") + ggtitle("Wild females") + theme(plot.title =
element_text(hjust = 0.5))+theme_classic()+geom_point(size=3)+
scale_color_manual(values = c("firebrick", "dodgerblue"))
ggplot(WildFemales, aes(x=RCBR_log, y=vLO_log, color=Subspecies)) +
geom_point()+labs(y= "Ventral lobulla [Log]", x = "Central Brain (minus AL & AOTu)
[Log]") + ggtitle("Wild females") + theme(plot.title = element_text(hjust = 0.5))
+theme_classic()+geom_point(size=3)+ scale_color_manual(values = c("firebrick",
"dodgerblue"))
ggplot(WildFemales, aes(x=RCBR_log, y=AME_log, color=Subspecies)) +
geom_point()+labs(y= "Accessory medulla [Log]", x = "Central Brain (minus AL &
AOTu) [Log]") + ggtitle("Wild females") + theme(plot.title = element_text(hjust = 0.5))
+theme_classic()+geom_point(size=3)+ scale_color_manual(values = c("firebrick",
"dodgerblue"))
ggplot(WildFemales, aes(x=RCBR_log, y=AOTu_log, color=Subspecies)) +
geom_point()+labs(y= "Anterior optic tubercle [Log]", x = "Central Brain (minus AL &
AOTu) [Log]") + ggtitle("Wild females") + theme(plot.title = element_text(hjust = 0.5))
+theme_classic()+geom_point(size=3)+ scale_color_manual(values = c("firebrick",
"dodgerblue"))
ggplot(WildFemales, aes(x=RCBR_log, y=AL_log, color=Subspecies)) +
geom_point()+labs(y= "Antennal lobe [Log]", x = "Central Brain (minus AL & AOTu)
[Log]") + ggtitle("Wild females") + theme(plot.title = element_text(hjust = 0.5))
+theme_classic()+geom_point(size=3)+ scale_color_manual(values = c("firebrick",
"dodgerblue"))
ggplot(WildFemales, aes(x=RCBR_log, y=LA_log, color=Subspecies)) +
geom_point()+labs(y= "Lamina [Log]", x = "Central Brain (minus AL & AOTu) [Log]")
+ ggtitle("Wild females") + theme(plot.title = element_text(hjust = 0.5))
+theme_classic()+geom_point(size=3)+ scale_color_manual(values = c("firebrick",
"dodgerblue"))

#Wild caught animals grouped as sexes and treated as a whole

medullawilderatoplot<-ggplot(WildBrains, aes(x=RCBR_log, y=ME_log, color=Sex))
+ geom_point()+labs(y= "Medulla [Log]", x = "Central Brain (minus AL & AOTu)
[Log]") + ggtitle("Wild caught") + theme(plot.title = element_text(hjust = 0.5))
+theme_classic()+geom_point(size=3)+ scale_color_manual(values = c("#00BC59",
"#E08B00"))
lobulawilderatoplot<-ggplot(WildBrains, aes(x=RCBR_log, y=LO_log, color=Sex)) +
geom_point()+labs(y= "Lobula [Log]", x = "Central Brain (minus AL & AOTu) [Log]") +
ggtitle("Wild caught") + theme(plot.title = element_text(hjust = 0.5))+theme_classic()
```

```
+geom_point(size=3)+ scale_color_manual(values = c("#00BC59", "#E08B00"))
lobulaplatewilderatoplot<-ggplot(WildBrains, aes(x=RCBR_log, y=LOP_log,
color=Sex)) + geom_point()+labs(y= "Lobula plate [Log]", x = "Central Brain (minus
AL & AOTu) [Log]") + ggtitle("Wild caught") + theme(plot.title = element_text(hjust =
0.5))+theme_classic()+geom_point(size=3)+ scale_color_manual(values =
c("#00BC59", "#E08B00"))
vlobulawilderatoplot<-ggplot(WildBrains, aes(x=RCBR_log, y=vLO_log, color=Sex))
+ geom_point()+labs(y= "Ventral lobulla [Log]", x = "Central Brain (minus AL & AOTu)
[Log]") + ggtitle("Wild caught") + theme(plot.title = element_text(hjust = 0.5))
+theme_classic()+geom_point(size=3)+ scale_color_manual(values = c("#00BC59",
"#E08B00"))
ggplot(WildBrains, aes(x=RCBR_log, y=AME_log, color=Sex)) + geom_point()
+labs(y= "Accessory medulla [Log]", x = "Central Brain (minus AL & AOTu) [Log]") +
ggtitle("Wild caught") + theme(plot.title = element_text(hjust = 0.5))+theme_classic()
+geom_point(size=3)+ scale_color_manual(values = c("#00BC59", "#E08B00"))
ggplot(WildBrains, aes(x=RCBR_log, y=AOTu_log, color=Sex)) + geom_point()
+labs(y= "Anterior optic tubercle [Log]", x = "Central Brain (minus AL & AOTu) [Log]")
+ ggtitle("Wild caught") + theme(plot.title = element_text(hjust = 0.5))
+theme_classic()+geom_point(size=3)+ scale_color_manual(values = c("#00BC59",
"#E08B00"))
alobewilderatoplot<-ggplot(WildBrains, aes(x=RCBR_log, y=AL_log, color=Sex)) +
geom_point()+labs(y= "Antennal lobe [Log]", x = "Central Brain (minus AL & AOTu)
[Log]") + ggtitle("Wild caught") + theme(plot.title = element_text(hjust = 0.5))
+theme_classic()+geom_point(size=3)+ scale_color_manual(values = c("#00BC59",
"#E08B00"))
laminawilderatoplot<-ggplot(WildBrains, aes(x=RCBR_log, y=LA_log, color=Sex)) +
geom_point()+labs(y= "Lamina [Log]", x = "Central Brain (minus AL & AOTu) [Log]")
+ ggtitle("Wild caught") + theme(plot.title = element_text(hjust = 0.5))
+theme_classic()+geom_point(size=3)+ scale_color_manual(values = c("#00BC59",
"#E08B00"))

#Wild caught chestertonii grouped as sexes

ggplot(Wildchestertonii, aes(x=RCBR_log, y=ME_log, color=Sex)) + geom_point()
+labs(y= "Medulla [Log]", x = "Central Brain (minus AL & AOTu) [Log]") +
ggtitle("Wild chestertonii") + theme(plot.title = element_text(hjust = 0.5))
+theme_classic()+geom_point(size=3)+ scale_color_manual(values = c("#00BC59",
"#E08B00"))
lobulawildchestertoniiplot<-ggplot(Wildchestertonii, aes(x=RCBR_log, y=LO_log,
color=Sex)) + geom_point()+labs(y= "Lobula [Log]", x = "Central Brain (minus AL &
AOTu) [Log]") + ggtitle("Wild chestertonii") + theme(plot.title = element_text(hjust =
0.5))+theme_classic()+geom_point(size=3)+ scale_color_manual(values =
c("#00BC59", "#E08B00"))
lobulaplatewildchestertoniiplot<-ggplot(Wildchestertonii, aes(x=RCBR_log,
y=LOP_log, color=Sex)) + geom_point()+labs(y= "Lobula plate [Log]", x = "Central
Brain (minus AL & AOTu) [Log]") + ggtitle("Wild chestertonii") + theme(plot.title =
element_text(hjust = 0.5))+theme_classic()+geom_point(size=3)+
scale_color_manual(values = c("#00BC59", "#E08B00"))
vlobulawildchestertoniiplot<-ggplot(Wildchestertonii, aes(x=RCBR_log, y=vLO_log,
color=Sex)) + geom_point()+labs(y= "Ventral lobula [Log]", x = "Central Brain (minus
```

```
AL & AOTu) [Log]") + ggtitle("Wild chestertonii") + theme(plot.title =
element_text(hjust = 0.5))+theme_classic()+geom_point(size=3)+
scale_color_manual(values = c("#00BC59", "#E08B00"))
ggplot(Wildchestertonii, aes(x=RCBR_log, y=AME_log, color=Sex)) + geom_point()
+labs(y= "Accessory medulla [Log]", x = "Central Brain (minus AL & AOTu) [Log]") +
ggtitle("Wild chestertonii") + theme(plot.title = element_text(hjust = 0.5))
+theme_classic()+geom_point(size=3)+ scale_color_manual(values = c("#00BC59",
"#E08B00"))
ggplot(Wildchestertonii, aes(x=RCBR_log, y=AOTu_log, color=Sex)) + geom_point()
+labs(y= "Anterior optic tubercle [Log]", x = "Central Brain (minus AL & AOTu) [Log]")
+ ggtitle("Wild chestertonii") + theme(plot.title = element_text(hjust = 0.5))
+theme_classic()+geom_point(size=3)+ scale_color_manual(values = c("#00BC59",
"#E08B00"))
ggplot(Wildchestertonii, aes(x=RCBR_log, y=AL_log, color=Sex)) + geom_point()
+labs(y= "Antennal lobe [Log]", x = "Central Brain (minus AL & AOTu) [Log]") +
ggtitle("Wild chestertonii") + theme(plot.title = element_text(hjust = 0.5))
+theme_classic()+geom_point(size=3)+ scale_color_manual(values = c("#00BC59",
"#E08B00"))
ggplot(Wildchestertonii, aes(x=RCBR_log, y=LA_log, color=Sex)) + geom_point()
+labs(y= "Lamina [Log]", x = "Central Brain (minus AL & AOTu) [Log]") + ggtitle("Wild
chestertonii") + theme(plot.title = element_text(hjust = 0.5))+theme_classic()
+geom_point(size=3)+ scale_color_manual(values = c("#00BC59", "#E08B00"))

#Wild caught venus grouped as sexes

medullawildvenusplot<-ggplot(Wildvenus, aes(x=RCBR_log, y=ME_log, color=Sex))
+ geom_point()+labs(y= "Medulla [Log]", x = "Central Brain (minus AL & AOTu)
[Log]") + ggtitle("Wild venus") + theme(plot.title = element_text(hjust = 0.5))
+theme_classic()+geom_point(size=3)+ scale_color_manual(values = c("#00BC59",
"#E08B00"))
ggplot(Wildvenus, aes(x=RCBR_log, y=LO_log, color=Sex)) + geom_point()+labs(y=
"Lobula [Log]", x = "Central Brain (minus AL & AOTu) [Log]") + ggtitle("Wild venus") +
theme(plot.title = element_text(hjust = 0.5))+theme_classic()+geom_point(size=3)+
scale_color_manual(values = c("#00BC59", "#E08B00"))
ggplot(Wildvenus, aes(x=RCBR_log, y=LOP_log, color=Sex)) + geom_point()
+labs(y= "Lobula plate [Log]", x = "Central Brain (minus AL & AOTu) [Log]") +
ggtitle("Wild venus") + theme(plot.title = element_text(hjust = 0.5))+theme_classic()
+geom_point(size=3)+ scale_color_manual(values = c("#00BC59", "#E08B00"))
ggplot(Wildvenus, aes(x=RCBR_log, y=vLO_log, color=Sex)) + geom_point()
+labs(y= "Ventral lobulla [Log]", x = "Central Brain (minus AL & AOTu) [Log]") +
ggtitle("Wild venus") + theme(plot.title = element_text(hjust = 0.5))+theme_classic()
+geom_point(size=3)+ scale_color_manual(values = c("#00BC59", "#E08B00"))
ggplot(Wildvenus, aes(x=RCBR_log, y=AME_log, color=Sex)) + geom_point()
+labs(y= "Accessory medulla [Log]", x = "Central Brain (minus AL & AOTu) [Log]") +
ggtitle("Wild venus") + theme(plot.title = element_text(hjust = 0.5))+theme_classic()
+geom_point(size=3)+ scale_color_manual(values = c("#00BC59", "#E08B00"))
ggplot(Wildvenus, aes(x=RCBR_log, y=AOTu_log, color=Sex)) + geom_point()
+labs(y= "Anterior optic tubercle [Log]", x = "Central Brain (minus AL & AOTu) [Log]")
+ ggtitle("Wild venus") + theme(plot.title = element_text(hjust = 0.5))
+theme_classic()+geom_point(size=3)+ scale_color_manual(values = c("#00BC59",
```

```
"#E08B00"))
alobewildvenusplot<-ggplot(Wildvenus, aes(x=RCBR_log, y=AL_log, color=Sex)) +
geom_point()+labs(y= "Antennal lobe [Log]", x = "Central Brain (minus AL & AOTu)
[Log]") + ggtitle("Wild venus") + theme(plot.title = element_text(hjust = 0.5))
+theme_classic()+geom_point(size=3)+ scale_color_manual(values = c("#00BC59",
"#E08B00"))
laminawildvenusplot<-ggplot(Wildvenus, aes(x=RCBR_log, y=LA_log, color=Sex)) +
geom_point()+labs(y= "Lamina [Log]", x = "Central Brain (minus AL & AOTu) [Log]")
+ ggtitle("Wild venus") + theme(plot.title = element_text(hjust = 0.5))
+theme_classic()+geom_point(size=3)+ scale_color_manual(values = c("#00BC59",
"#E08B00"))

#Analyses in insectary samples

RCBRreducedi<-lm(RCBR_log ~Subspecies+Sex, data = InsectaryBrains)
RCBRalti<-lm(RCBR_log ~Subspecies, data = InsectaryBrains)
RCBRalt1i<-lm(RCBR_log ~Sex, data = InsectaryBrains)

anova(RCBRreducedi,RCBRalti)


summary(RCBRreducedw)
summary(RCBRalti)
summary(RCBRalt1i)



#Medulla Modelling (Subspecies*)


MedullaFull<-lmer(ME_log ~ RCBR_log *Subspecies +(1|Sex), data =
InsectaryBrains)

drop1(MedullaFull, test="Chisq")
newmedulla<-update(MedullaFull, ~. -RCBR_log:Subspecies)

drop1(newmedulla, test="Chisq")

MedullaReduced<-lmer(ME_log ~ RCBR_log +Subspecies+(1|Sex), data =
InsectaryBrains)
MedullaAlt<-lmer(ME_log ~ RCBR_log+(1|Sex), data = InsectaryBrains)

anova(MedullaReduced, MedullaAlt)
Anova(MedullaReduced)

summary(MedullaReduced)

#Lobulla Modelling (Subspecies*)
```

```
LobulaFull<-lmer(LO_log ~ RCBR_log *Subspecies +(1|Sex), data =
InsectaryBrains)

drop1(LobulaFull, test="Chisq")
newlobula<-update(LobulaFull, ~. -RCBR_log:Subspecies)

drop1(newlobula, test="Chisq")


LobulaReduced<-lmer(LO_log ~ RCBR_log +Subspecies+(1|Sex), data =
InsectaryBrains)
LobulaAlt<-lmer(LO_log ~ RCBR_log+(1|Sex), data = InsectaryBrains)

anova(LobulaReduced, LobulaAlt)
Anova(LobulaReduced)

summary(LobulaReduced)



#Lobulla plate models (Subspecies*)

LobulapFull<-lmer(LOP_log ~ RCBR_log *Subspecies +(1|Sex), data =
InsectaryBrains)

drop1(LobulapFull, test="Chisq")
newlobulap<-update(LobulaFull, ~. -RCBR_log:Subspecies)

drop1(newlobulap, test="Chisq")


LobulaPReduced<-lmer(LOP_log ~ RCBR_log +Subspecies+(1|Sex), data =
InsectaryBrains)
LobulapAlt<-lmer(LOP_log ~ RCBR_log+(1|Sex), data = InsectaryBrains)

anova(LobulaPReduced, LobulapAlt)
Anova(LobulaPReduced)

summary(LobulaPReduced)

#Ventral lobula modelling (Subspecies*)


vLobulaFull<-lmer(vLO_log ~ RCBR_log *Subspecies +(1|Sex), data =
InsectaryBrains)

drop1(vLobulaFull, test="Chisq")
newvlobula<-update(vLobulaFull, ~. -RCBR_log:Subspecies)

drop1(newvlobula, test="Chisq")
```

```
vLobulaReduced<-lmer(vLO_log ~ RCBR_log+Subspecies+(1|Sex), data =
InsectaryBrains)
vLobulaAlt<-lmer(vLO_log ~ RCBR_log+(1|Sex), data = InsectaryBrains)

anova(vLobulaReduced, vLobulaAlt)
Anova(vLobulaReduced)

summary(vLobulaReduced)



#Lamina modelling (Subspecies*)

LaminaFull<-lmer(LA_log ~ RCBR_log *Subspecies+(1|Sex) , data =
InsectaryBrains)

drop1(LaminaFull, test="Chisq")
newlamina<-update(LaminaFull, ~. -RCBR_log:Subspecies)

drop1(newlamina, test="Chisq")

LaminaReduced<-lmer(LA_log ~ RCBR_log+Subspecies+(1|Sex), data =
InsectaryBrains)
LaminaAlt<-lmer(LA_log ~ RCBR_log +(1|Sex), data = InsectaryBrains)

anova(LaminaReduced, LaminaAlt)
Anova(LaminaReduced)


summary(LaminaReduced)



#Accessory medulla modelling ()

ameFull<-lmer(AME_log ~ RCBR_log *Subspecies +(1|Sex), data = InsectaryBrains)

drop1(ameFull, test="Chisq")
newame<-update(ameFull, ~. -RCBR_log:Subspecies)

drop1(newame, test="Chisq")
newame1<-update(newame, ~. -Subspecies)

drop1(newame1, test="Chisq")


ameReduced<-lmer(AME_log ~ RCBR_log+(1|Sex), data = InsectaryBrains)
ameAlt<-lmer(AME_log ~ RCBR_log+Subspecies+(1|Sex), data = InsectaryBrains)
```

```
anova(ameReduced, ameAlt)
Anova(ameAlt)

summary(ameReduced)




#Anterior Optic Tubercle (Subspecies*)

AOTuFull<-lmer(AOTu_log ~ RCBR_log *Subspecies +(1|Sex), data =
InsectaryBrains)

drop1(AOTuFull, test="Chisq")
newAOTu<-update(AOTuFull, ~. -RCBR_log:Subspecies)

drop1(newAOTu, test="Chisq")


AOTureduced<-lmer(AOTu_log ~ RCBR_log+Subspecies+(1|Sex), data =
InsectaryBrains)
AOTuAlt<-lmer(AOTu_log ~ RCBR_log+(1|Sex), data = InsectaryBrains)

anova(AOTureduced, AOTuAlt)
Anova(AOTureduced)

summary(AOTureduced)



#Antennal Lobe models (Subspecies*)


alobeFull<-lmer(AL_log ~ RCBR_log *Subspecies +(1|Sex), data = InsectaryBrains)

drop1(alobeFull, test="Chisq")
alobenew<-update(alobeFull, ~. -RCBR_log:Subspecies)

drop1(alobenew, test="Chisq")


alobeReduced<-lmer(AL_log ~ RCBR_log+Subspecies+(1|Sex), data =
InsectaryBrains)
alobeAlt<-lmer(AL_log ~ RCBR_log+(1|Sex), data = InsectaryBrains)

anova(alobeReduced, alobeAlt)
Anova(alobeReduced)

summary(alobeReduced)
```

```
#Estimating centroids for the lobula plate of insectary reared butterflies


centroid_loburcbr_chester<- mean(Insectarychestertonii$RCBR_log)
centroid_lobulap_chester<- mean(Insectarychestertonii$LOP_log)


centroid_loburcbr_venus<- mean(Insectaryvenus$RCBR_log)
centroid_lobulap_venus<- mean(Insectaryvenus$LOP_log)


#Exploratory plots (just quickly checking the scaling relationships are not too off)

#Insectary reared animals grouped as species

medullainsectplot<-ggplot(InsectaryBrains, aes(x=RCBR_log, y=ME_log,
color=Subspecies)) + geom_point()+labs(x = bquote('[Log] rCBR'~(µm^3)), y =
bquote('[Log] Medulla'~(µm^3))) + ggtitle("Insectary reared") + theme(plot.title =
element_text(hjust = 0.5))+theme_classic()+geom_point(size=3)+
scale_color_manual(values = c("red", "dodgerblue"))+theme(axis.title =
element_text(size = 15))
lobulainsectplot<-ggplot(InsectaryBrains, aes(x=RCBR_log, y=LO_log,
color=Subspecies)) + geom_point()+labs(x = bquote('[Log] rCBR'~(µm^3)), y =
bquote('[Log] Lobula'~(µm^3))) + ggtitle("Insectary reared") + theme(plot.title =
element_text(hjust = 0.5))+theme_classic()+geom_point(size=3)+
scale_color_manual(values = c("red", "dodgerblue"))+theme(axis.title =
element_text(size = 15))
lopbulaplateinsectplot<-ggplot(InsectaryBrains, aes(x=RCBR_log, y=LOP_log,
color=Subspecies)) + geom_point()+labs(x = bquote('[Log] rCBR'~(µm^3)), y =
bquote('[Log] Lobula plate'~(µm^3))) + ggtitle("Insectary reared") + theme(plot.title =
element_text(hjust = 0.5))+theme_classic()+geom_point(size=3)+
scale_color_manual(values = c("red", "dodgerblue"))+theme(axis.title =
element_text(size = 15))+ annotate("point",x=8.071798,y=6.82714, col="red",
cex=11, shape=3)+annotate("point",x=8.11742,y=6.898365, col="blue", cex=11,
shape=3)
vlobulainsectplot<-ggplot(InsectaryBrains, aes(x=RCBR_log, y=vLO_log,
color=Subspecies)) + geom_point()+labs(y= "Ventral lobulla [Log]", x = "Rest of the
Central Brain [Log]") + ggtitle("Insectary reared") + theme(plot.title =
element_text(hjust = 0.5))+theme_classic()+geom_point(size=3)+
scale_color_manual(values = c("red", "dodgerblue"))+theme(axis.title =
element_text(size = 15))
amedullainsectplot<-ggplot(InsectaryBrains, aes(x=RCBR_log, y=AME_log,
color=Subspecies)) + geom_point()+labs(y= "Accessory medulla [Log]", x = "Rest of
the Central Brain [Log]") + ggtitle("Insectary reared") + theme(plot.title =
element_text(hjust = 0.5))+theme_classic()+geom_point(size=3)+
scale_color_manual(values = c("red", "dodgerblue"))+theme(axis.title =
element_text(size = 15))
aotubercleinsectplot<-ggplot(InsectaryBrains, aes(x=RCBR_log, y=AOTu_log,
```

```
color=Subspecies)) + geom_point()+labs(y= "Anterior optic tubercle [Log]", x = "Rest of the Central Brain [Log]") + ggtitle("Insectary reared") + theme(plot.title = element_text(hjust = 0.5))+theme_classic()+geom_point(size=3)+ scale_color_manual(values = c("red", "dodgerblue"))+theme(axis.title = element_text(size = 15))
alobeinsectplot<-ggplot(InsectaryBrains, aes(x=RCBR_log, y=AL_log, color=Subspecies)) + geom_point()+labs(x = bquote('[Log] rCBR'~(µm^3)), y = bquote('[Log] Antennal lobe'~(µm^3))) + ggtitle("Insectary reared") + theme(plot.title = element_text(hjust = 0.5))+theme_classic()+geom_point(size=3)+ scale_color_manual(values = c("red", "dodgerblue"))+theme(axis.title = element_text(size = 15))
laminainsectplot<-ggplot(InsectaryBrains, aes(x=RCBR_log, y=LA_log, color=Subspecies)) + geom_point()+labs(y= "Lamina [Log]", x = "Rest of the Central Brain [Log]") + ggtitle("Insectary reared") + theme(plot.title = element_text(hjust = 0.5))+theme_classic()+geom_point(size=3)+ scale_color_manual(values = c("red", "dodgerblue"))+theme(axis.title = element_text(size = 15))
potuberclewildplot<-ggplot(InsectaryBrains, aes(x=RCBR_log, y=POTu_log, color=Subspecies)) + geom_point()+labs(y= "Posterior optic tubercle [Log]", x = "Rest of the Central Brain [Log]") + ggtitle("Insectary reared") + theme(plot.title = element_text(hjust = 0.5))+theme_classic()+geom_point(size=3)+ scale_color_manual(values = c("red", "dodgerblue"))+theme(axis.title = element_text(size = 15))

#Insectary reared males grouped as species

ggplot(InsectaryMales, aes(x=RCBR_log, y=ME_log, color=Subspecies)) + geom_point()+labs(y= "Medulla [Log]", x = "Central Brain (minus AL & AOTu) [Log]") + ggtitle("Insectary reared males") + theme(plot.title = element_text(hjust = 0.5)) +theme_classic()+geom_point(size=3)+ scale_color_manual(values = c("firebrick", "dodgerblue"))
ggplot(InsectaryMales, aes(x=RCBR_log, y=LO_log, color=Subspecies)) + geom_point()+labs(y= "Lobula [Log]", x = "Central Brain (minus AL & AOTu) [Log]") + ggtitle("Insectary reared males") + theme(plot.title = element_text(hjust = 0.5)) +theme_classic()+geom_point(size=3)+ scale_color_manual(values = c("firebrick", "dodgerblue"))
ggplot(InsectaryMales, aes(x=RCBR_log, y=LOP_log, color=Subspecies)) + geom_point()+labs(y= "Lobula plate [Log]", x = "Central Brain (minus AL & AOTu) [Log]") + ggtitle("Insectary reared males") + theme(plot.title = element_text(hjust = 0.5))+theme_classic()+geom_point(size=3)+ scale_color_manual(values = c("firebrick", "dodgerblue"))
ggplot(InsectaryMales, aes(x=RCBR_log, y=vLO_log, color=Subspecies)) + geom_point()+labs(y= "Ventral lobulla [Log]", x = "Central Brain (minus AL & AOTu) [Log]") + ggtitle("Insectary reared males") + theme(plot.title = element_text(hjust = 0.5))+theme_classic()+geom_point(size=3)+ scale_color_manual(values = c("firebrick", "dodgerblue"))
ggplot(InsectaryMales, aes(x=RCBR_log, y=AME_log, color=Subspecies)) + geom_point()+labs(y= "Accessory medulla [Log]", x = "Central Brain (minus AL & AOTu) [Log]") + ggtitle("Insectary reared males") + theme(plot.title = element_text(hjust = 0.5))+theme_classic()+geom_point(size=3)+ scale_color_manual(values = c("firebrick", "dodgerblue"))
```

```
ggplot(InsectaryMales, aes(x=RCBR_log, y=AOTu_log, color=Subspecies)) + geom_point()+labs(y= "Anterior optic tubercle [Log]", x = "Central Brain (minus AL & AOTu) [Log]") + ggtitle("Insectary reared males") + theme(plot.title = element_text(hjust = 0.5))+theme_classic()+geom_point(size=3)+ scale_color_manual(values = c("firebrick", "dodgerblue"))
ggplot(InsectaryMales, aes(x=RCBR_log, y=AL_log, color=Subspecies)) + geom_point()+labs(y= "Antennal lobe [Log]", x = "Central Brain (minus AL & AOTu) [Log]") + ggtitle("Insectary reared males") + theme(plot.title = element_text(hjust = 0.5))+theme_classic()+geom_point(size=3)+ scale_color_manual(values = c("firebrick", "dodgerblue"))
ggplot(InsectaryMales, aes(x=RCBR_log, y=LA_log, color=Subspecies)) + geom_point()+labs(y= "Lamina [Log]", x = "Central Brain (minus AL & AOTu) [Log]") + ggtitle("Insectary reared males") + theme(plot.title = element_text(hjust = 0.5)) +theme_classic()+geom_point(size=3)+ scale_color_manual(values = c("firebrick", "dodgerblue"))

#Insectary reared females separated by species

ggplot(InsectaryFemales, aes(x=RCBR_log, y=ME_log, color=Subspecies)) + geom_point()+labs(y= "Medulla [Log]", x = "Central Brain (minus AL & AOTu) [Log]") + ggtitle("Insectary reared females") + theme(plot.title = element_text(hjust = 0.5)) +theme_classic()+geom_point(size=3)+ scale_color_manual(values = c("firebrick", "dodgerblue"))
ggplot(InsectaryFemales, aes(x=RCBR_log, y=LO_log, color=Subspecies)) + geom_point()+labs(y= "Lobula [Log]", x = "Central Brain (minus AL & AOTu) [Log]") + ggtitle("Insectary reared females") + theme(plot.title = element_text(hjust = 0.5)) +theme_classic()+geom_point(size=3)+ scale_color_manual(values = c("firebrick", "dodgerblue"))
ggplot(InsectaryFemales, aes(x=RCBR_log, y=LOP_log, color=Subspecies)) + geom_point()+labs(y= "Lobula plate [Log]", x = "Central Brain (minus AL & AOTu) [Log]") + ggtitle("Insectary reared females") + theme(plot.title = element_text(hjust = 0.5))+theme_classic()+geom_point(size=3)+ scale_color_manual(values = c("firebrick", "dodgerblue"))
ggplot(InsectaryFemales, aes(x=RCBR_log, y=vLO_log, color=Subspecies)) + geom_point()+labs(y= "Ventral lobulla [Log]", x = "Central Brain (minus AL & AOTu) [Log]") + ggtitle("Insectary reared females") + theme(plot.title = element_text(hjust = 0.5))+theme_classic()+geom_point(size=3)+ scale_color_manual(values = c("firebrick", "dodgerblue"))
ggplot(InsectaryFemales, aes(x=RCBR_log, y=AME_log, color=Subspecies)) + geom_point()+labs(y= "Accessory medulla [Log]", x = "Central Brain (minus AL & AOTu) [Log]") + ggtitle("Insectary reared females") + theme(plot.title = element_text(hjust = 0.5))+theme_classic()+geom_point(size=3)+ scale_color_manual(values = c("firebrick", "dodgerblue"))
ggplot(InsectaryFemales, aes(x=RCBR_log, y=AOTu_log, color=Subspecies)) + geom_point()+labs(y= "Anterior optic tubercle [Log]", x = "Central Brain (minus AL & AOTu) [Log]") + ggtitle("Insectary reared females") + theme(plot.title = element_text(hjust = 0.5))+theme_classic()+geom_point(size=3)+ scale_color_manual(values = c("firebrick", "dodgerblue"))
ggplot(InsectaryFemales, aes(x=RCBR_log, y=AL_log, color=Subspecies)) + geom_point()+labs(y= "Antennal lobe [Log]", x = "Central Brain (minus AL & AOTu)
```

```
[Log]") + ggtitle("Insectary reared females") + theme(plot.title = element_text(hjust =
0.5))+theme_classic()+geom_point(size=3)+ scale_color_manual(values =
c("firebrick", "dodgerblue"))
ggplot(InsectaryFemales, aes(x=RCBR_log, y=LA_log, color=Subspecies)) +
geom_point()+labs(y= "Lamina [Log]", x = "Central Brain (minus AL & AOTu) [Log]")
+ ggtitle("Insectary reared females") + theme(plot.title = element_text(hjust = 0.5))
+theme_classic()+geom_point(size=3)+ scale_color_manual(values = c("firebrick",
"dodgerblue"))

#Insectary reared animals grouped as sexes and treated as a whole

medullainsectplot<-ggplot(InsectaryBrains, aes(x=RCBR_log, y=ME_log,
color=Sex)) + geom_point()+labs(y= "Medulla [Log]", x = "Central Brain (minus AL &
AOTu) [Log]") + ggtitle("Insectary reared") + theme(plot.title = element_text(hjust =
0.5))+theme_classic()+geom_point(size=3)+ scale_color_manual(values =
c("#00BC59", "#E08B00"))
ggplot(InsectaryBrains, aes(x=RCBR_log, y=LO_log, color=Sex)) + geom_point()
+labs(y= "Lobula [Log]", x = "Central Brain (minus AL & AOTu) [Log]") +
ggtitle("Insectary reared") + theme(plot.title = element_text(hjust = 0.5))
+theme_classic()+geom_point(size=3)+ scale_color_manual(values = c("#00BC59",
"#E08B00"))
ggplot(InsectaryBrains, aes(x=RCBR_log, y=LOP_log, color=Sex)) + geom_point()
+labs(y= "Lobula plate [Log]", x = "Central Brain (minus AL & AOTu) [Log]") +
ggtitle("Insectary reared") + theme(plot.title = element_text(hjust = 0.5))
+theme_classic()+geom_point(size=3)+ scale_color_manual(values = c("#00BC59",
"#E08B00"))
ggplot(InsectaryBrains, aes(x=RCBR_log, y=vLO_log, color=Sex)) + geom_point()
+labs(y= "Ventral lobulla [Log]", x = "Central Brain (minus AL & AOTu) [Log]") +
ggtitle("Insectary reared") + theme(plot.title = element_text(hjust = 0.5))
+theme_classic()+geom_point(size=3)+ scale_color_manual(values = c("#00BC59",
"#E08B00"))
ggplot(InsectaryBrains, aes(x=RCBR_log, y=AME_log, color=Sex)) + geom_point()
+labs(y= "Accessory medulla [Log]", x = "Central Brain (minus AL & AOTu) [Log]") +
ggtitle("Insectary reared") + theme(plot.title = element_text(hjust = 0.5))
+theme_classic()+geom_point(size=3)+ scale_color_manual(values = c("#00BC59",
"#E08B00"))
ggplot(InsectaryBrains, aes(x=RCBR_log, y=AOTu_log, color=Sex)) + geom_point()
+labs(y= "Anterior optic tubercle [Log]", x = "Central Brain (minus AL & AOTu) [Log]")
+ ggtitle("Insectary reared") + theme(plot.title = element_text(hjust = 0.5))
+theme_classic()+geom_point(size=3)+ scale_color_manual(values = c("#00BC59",
"#E08B00"))
ggplot(InsectaryBrains, aes(x=RCBR_log, y=AL_log, color=Sex)) + geom_point()
+labs(y= "Antennal lobe [Log]", x = "Central Brain (minus AL & AOTu) [Log]") +
ggtitle("Insectary reared") + theme(plot.title = element_text(hjust = 0.5))
+theme_classic()+geom_point(size=3)+ scale_color_manual(values = c("#00BC59",
"#E08B00"))
laminainsectplot<-ggplot(InsectaryBrains, aes(x=RCBR_log, y=LA_log, color=Sex))
+ geom_point()+labs(y= "Lamina [Log]", x = "Central Brain (minus AL & AOTu)
[Log]") + ggtitle("Insectary reared") + theme(plot.title = element_text(hjust = 0.5))
+theme_classic()+geom_point(size=3)+ scale_color_manual(values = c("#00BC59",
```

```
"#E08B00"))

#Insectary reared chestertonii grouped as sexes

ggplot(Insectarychestertonii, aes(x=RCBR_log, y=ME_log, color=Sex)) +
geom_point()+labs(y= "Medulla [Log]", x = "Central Brain (minus AL & AOTu) [Log]")
+ ggtitle("Insectary reared chestertonii") + theme(plot.title = element_text(hjust =
0.5))+theme_classic()+geom_point(size=3)+ scale_color_manual(values =
c("#00BC59", "#E08B00"))
ggplot(Insectarychestertonii, aes(x=RCBR_log, y=LO_log, color=Sex)) +
geom_point()+labs(y= "Lobula [Log]", x = "Central Brain (minus AL & AOTu) [Log]") +
ggtitle("Insectary reared chestertonii") + theme(plot.title = element_text(hjust = 0.5))
+theme_classic()+geom_point(size=3)+ scale_color_manual(values = c("#00BC59",
"#E08B00"))
ggplot(Insectarychestertonii, aes(x=RCBR_log, y=LOP_log, color=Sex)) +
geom_point()+labs(y= "Lobula plate [Log]", x = "Central Brain (minus AL & AOTu)
[Log]") + ggtitle("Insectary reared chestertonii") + theme(plot.title =
element_text(hjust = 0.5))+theme_classic()+geom_point(size=3)+
scale_color_manual(values = c("#00BC59", "#E08B00"))
ggplot(Insectarychestertonii, aes(x=RCBR_log, y=vLO_log, color=Sex)) +
geom_point()+labs(y= "Ventral lobula [Log]", x = "Central Brain (minus AL & AOTu)
[Log]") + ggtitle("Insectary reared chestertonii") + theme(plot.title =
element_text(hjust = 0.5))+theme_classic()+geom_point(size=3)+
scale_color_manual(values = c("#00BC59", "#E08B00"))
ggplot(Insectarychestertonii, aes(x=RCBR_log, y=AME_log, color=Sex)) +
geom_point()+labs(y= "Accessory medulla [Log]", x = "Central Brain (minus AL &
AOTu) [Log]") + ggtitle("Insectary reared chestertonii") + theme(plot.title =
element_text(hjust = 0.5))+theme_classic()+geom_point(size=3)+
scale_color_manual(values = c("#00BC59", "#E08B00"))
ggplot(Insectarychestertonii, aes(x=RCBR_log, y=AOTu_log, color=Sex)) +
geom_point()+labs(y= "Anterior optic tubercle [Log]", x = "Central Brain (minus AL &
AOTu) [Log]") + ggtitle("Insectary reared chestertonii") + theme(plot.title =
element_text(hjust = 0.5))+theme_classic()+geom_point(size=3)+
scale_color_manual(values = c("#00BC59", "#E08B00"))
ggplot(Insectarychestertonii, aes(x=RCBR_log, y=AL_log, color=Sex)) +
geom_point()+labs(y= "Antennal lobe [Log]", x = "Central Brain (minus AL & AOTu)
[Log]") + ggtitle("Insectary reared chestertonii") + theme(plot.title =
element_text(hjust = 0.5))+theme_classic()+geom_point(size=3)+
scale_color_manual(values = c("#00BC59", "#E08B00"))
ggplot(Insectarychestertonii, aes(x=RCBR_log, y=LA_log, color=Sex)) +
geom_point()+labs(y= "Lamina [Log]", x = "Central Brain (minus AL & AOTu) [Log]")
+ ggtitle("Insectary reared chestertonii") + theme(plot.title = element_text(hjust =
0.5))+theme_classic()+geom_point(size=3)+ scale_color_manual(values =
c("#00BC59", "#E08B00"))

#Insectary reared venus grouped as sexes

ggplot(Insectaryvenus, aes(x=RCBR_log, y=ME_log, color=Sex)) + geom_point()
+labs(y= "Medulla [Log]", x = "Central Brain (minus AL & AOTu) [Log]") +
ggtitle("Insectary reared venus") + theme(plot.title = element_text(hjust = 0.5))
```

```
+theme_classic()+geom_point(size=3)+ scale_color_manual(values = c("#00BC59", "#E08B00"))
ggplot(Insectaryvenus, aes(x=RCBR_log, y=LO_log, color=Sex)) + geom_point()
+labs(y= "Lobula [Log]", x = "Central Brain (minus AL & AOTu) [Log]") +
ggtitle("Insectary reared venus") + theme(plot.title = element_text(hjust = 0.5))
+theme_classic()+geom_point(size=3)+ scale_color_manual(values = c("#00BC59", "#E08B00"))
ggplot(Insectaryvenus, aes(x=RCBR_log, y=LOP_log, color=Sex)) + geom_point()
+labs(y= "Lobula plate [Log]", x = "Central Brain (minus AL & AOTu) [Log]") +
ggtitle("Insectary reared venus") + theme(plot.title = element_text(hjust = 0.5))
+theme_classic()+geom_point(size=3)+ scale_color_manual(values = c("#00BC59", "#E08B00"))
ggplot(Insectaryvenus, aes(x=RCBR_log, y=vLO_log, color=Sex)) + geom_point()
+labs(y= "Ventral lobulla [Log]", x = "Central Brain (minus AL & AOTu) [Log]") +
ggtitle("Insectary reared venus") + theme(plot.title = element_text(hjust = 0.5))
+theme_classic()+geom_point(size=3)+ scale_color_manual(values = c("#00BC59", "#E08B00"))
ggplot(Insectaryvenus, aes(x=RCBR_log, y=AME_log, color=Sex)) + geom_point()
+labs(y= "Accessory medulla [Log]", x = "Central Brain (minus AL & AOTu) [Log]") +
ggtitle("Insectary reared venus") + theme(plot.title = element_text(hjust = 0.5))
+theme_classic()+geom_point(size=3)+ scale_color_manual(values = c("#00BC59", "#E08B00"))
aotuinsectvenusplot<-ggplot(Insectaryvenus, aes(x=RCBR_log, y=AOTu_log, color=Sex)) + geom_point()+labs(y= "Anterior optic tubercle [Log]", x = "Central Brain (minus AL & AOTu) [Log]") + ggtitle("Insectary reared venus") + theme(plot.title = element_text(hjust = 0.5))+theme_classic()+geom_point(size=3)+ scale_color_manual(values = c("#00BC59", "#E08B00"))
alobeinsectvenusplot<-ggplot(Insectaryvenus, aes(x=RCBR_log, y=AL_log, color=Sex)) + geom_point()+labs(y= "Antennal lobe [Log]", x = "Central Brain (minus AL & AOTu) [Log]") + ggtitle("Insectary reared venus") + theme(plot.title = element_text(hjust = 0.5))+theme_classic()+geom_point(size=3)+ scale_color_manual(values = c("#00BC59", "#E08B00"))
ggplot(Insectaryvenus, aes(x=RCBR_log, y=LA_log, color=Sex)) + geom_point()
+labs(y= "Lamina [Log]", x = "Central Brain (minus AL & AOTu) [Log]") +
ggtitle("Insectary reared venus") + theme(plot.title = element_text(hjust = 0.5))
+theme_classic()+geom_point(size=3)+ scale_color_manual(values = c("#00BC59", "#E08B00"))

#All venus grouped by origin

ggplot(VenusBrains, aes(x=RCBR_log, y=ME_log, color=Origin)) + geom_point()
+labs(y= "Medulla [Log]", x = "Central Brain (minus AL & AOTu) [Log]") +
ggtitle("Insectary reared venus") + theme(plot.title = element_text(hjust = 0.5))
+theme_classic()+geom_point(size=3)+ scale_color_manual(values = c("#00BC59", "#E08B00"))
ggplot(VenusBrains, aes(x=RCBR_log, y=LO_log, color=Origin)) + geom_point()
+labs(y= "Lobula [Log]", x = "Central Brain (minus AL & AOTu) [Log]") +
ggtitle("Insectary reared venus") + theme(plot.title = element_text(hjust = 0.5))
+theme_classic()+geom_point(size=3)+ scale_color_manual(values = c("#00BC59", "#E08B00"))
```

```
ggplot(VenusBrains, aes(x=RCBR_log, y=LOP_log, color=Origin)) + geom_point()
+labs(y= "Lobula plate [Log]", x = "Central Brain (minus AL & AOTu) [Log]") +
ggtitle("Insectary reared venus") + theme(plot.title = element_text(hjust = 0.5))
+theme_classic()+geom_point(size=3)+ scale_color_manual(values = c("#00BC59",
"#E08B00"))
ggplot(VenusBrains, aes(x=RCBR_log, y=vLO_log, color=Origin)) + geom_point()
+labs(y= "Ventral lobula [Log]", x = "Central Brain (minus AL & AOTu) [Log]") +
ggtitle("Insectary reared venus") + theme(plot.title = element_text(hjust = 0.5))
+theme_classic()+geom_point(size=3)+ scale_color_manual(values = c("#00BC59",
"#E08B00"))
ggplot(VenusBrains, aes(x=RCBR_log, y=AME_log, color=Origin)) + geom_point()
+labs(y= "Accessory medulla [Log]", x = "Central Brain (minus AL & AOTu) [Log]") +
ggtitle("Insectary reared venus") + theme(plot.title = element_text(hjust = 0.5))
+theme_classic()+geom_point(size=3)+ scale_color_manual(values = c("#00BC59",
"#E08B00"))
ggplot(VenusBrains, aes(x=RCBR_log, y=AOTu_log, color=Origin)) + geom_point()
+labs(y= "Anterior optic tubercle [Log]", x = "Central Brain (minus AL & AOTu) [Log]")
+ ggtitle("Insectary reared venus") + theme(plot.title = element_text(hjust = 0.5))
+theme_classic()+geom_point(size=3)+ scale_color_manual(values = c("#00BC59",
"#E08B00"))
ggplot(VenusBrains, aes(x=RCBR_log, y=AL_log, color=Origin)) + geom_point()
+labs(y= "Antennal lobe [Log]", x = "Central Brain (minus AL & AOTu) [Log]") +
ggtitle("Insectary reared venus") + theme(plot.title = element_text(hjust = 0.5))
+theme_classic()+geom_point(size=3)+ scale_color_manual(values = c("#00BC59",
"#E08B00"))
ggplot(VenusBrains, aes(x=RCBR_log, y=LA_log, color=Origin)) + geom_point()
+labs(y= "Lamina [Log]", x = "Central Brain (minus AL & AOTu) [Log]") +
ggtitle("Insectary reared venus") + theme(plot.title = element_text(hjust = 0.5))
+theme_classic()+geom_point(size=3)+ scale_color_manual(values = c("#00BC59",
"#E08B00"))


#All chestertonii grouped by origin

ggplot(Chestertoniibrains, aes(x=RCBR_log, y=ME_log, color=Origin)) +
geom_point()+labs(y= "Medulla [Log]", x = "Central Brain (minus AL & AOTu) [Log]")
+ ggtitle("Insectary reared venus") + theme(plot.title = element_text(hjust = 0.5))
+theme_classic()+geom_point(size=3)+ scale_color_manual(values = c("#00BC59",
"#E08B00"))
ggplot(Chestertoniibrains, aes(x=RCBR_log, y=LO_log, color=Origin)) +
geom_point()+labs(y= "Lobula [Log]", x = "Central Brain (minus AL & AOTu) [Log]") +
ggtitle("Insectary reared venus") + theme(plot.title = element_text(hjust = 0.5))
+theme_classic()+geom_point(size=3)+ scale_color_manual(values = c("#00BC59",
"#E08B00"))
ggplot(Chestertoniibrains, aes(x=RCBR_log, y=LOP_log, color=Origin)) +
geom_point()+labs(y= "Lobula plate [Log]", x = "Central Brain (minus AL & AOTu)
[Log]") + ggtitle("Insectary reared venus") + theme(plot.title = element_text(hjust =
0.5))+theme_classic()+geom_point(size=3)+ scale_color_manual(values =
c("#00BC59", "#E08B00"))
ggplot(Chestertoniibrains, aes(x=RCBR_log, y=vLO_log, color=Origin)) +
```

```
geom_point()+labs(y= "Ventral lobula [Log]", x = "Central Brain (minus AL & AOTu) [Log]") + ggtitle("Insectary reared venus") + theme(plot.title = element_text(hjust = 0.5))+theme_classic()+geom_point(size=3)+ scale_color_manual(values = c("#00BC59", "#E08B00"))
ggplot(Chestertoniibrains, aes(x=RCBR_log, y=AME_log, color=Origin)) + geom_point()+labs(y= "Accessory medulla [Log]", x = "Central Brain (minus AL & AOTu) [Log]") + ggtitle("Insectary reared venus") + theme(plot.title = element_text(hjust = 0.5))+theme_classic()+geom_point(size=3)+ scale_color_manual(values = c("#00BC59", "#E08B00"))
ggplot(Chestertoniibrains, aes(x=RCBR_log, y=AOTu_log, color=Origin)) + geom_point()+labs(y= "Anterior optic tubercle [Log]", x = "Central Brain (minus AL & AOTu) [Log]") + ggtitle("Insectary reared venus") + theme(plot.title = element_text(hjust = 0.5))+theme_classic()+geom_point(size=3)+ scale_color_manual(values = c("#00BC59", "#E08B00"))
ggplot(Chestertoniibrains, aes(x=RCBR_log, y=AL_log, color=Origin)) + geom_point()+labs(y= "Antennal lobe [Log]", x = "Central Brain (minus AL & AOTu) [Log]") + ggtitle("Insectary reared venus") + theme(plot.title = element_text(hjust = 0.5))+theme_classic()+geom_point(size=3)+ scale_color_manual(values = c("#00BC59", "#E08B00"))
ggplot(Chestertoniibrains, aes(x=RCBR_log, y=LA_log, color=Origin)) + geom_point()+labs(y= "Lamina [Log]", x = "Central Brain (minus AL & AOTu) [Log]") + ggtitle("Insectary reared venus") + theme(plot.title = element_text(hjust = 0.5)) +theme_classic()+geom_point(size=3)+ scale_color_manual(values = c("#00BC59", "#E08B00"))



#SMATR on wild individuals grouped in subspecies

#Testing for correlation between the allometric control and the neuropil of interest (in all the cases it exists)

sma(WildBrains$ME_log~WildBrains$RCBR_log, robust = T)
sma(WildBrains$LO_log~WildBrains$RCBR_log, robust = T)
sma(WildBrains$LOP_log~WildBrains$RCBR_log, robust = T)
sma(WildBrains$vLO_log~WildBrains$RCBR_log, robust = T)
sma(WildBrains$AME_log~WildBrains$RCBR_log, robust = T)
sma(WildBrains$AOTu_log~WildBrains$RCBR_log, robust = T)
sma(WildBrains$AL_log~WildBrains$RCBR_log, robust = T)
sma(WildBrains$LA_log~WildBrains$RCBR_log, robust = T)
sma(WildBrains$POTu_log~WildBrains$RCBR_log, robust = T)


#Testing for slope (β) changes between subspecies (in all the neuropils, the slopes are similar)

sma(WildBrains$ME_log~WildBrains$RCBR_log*WildBrains$Subspecies, robust = T)
sma(WildBrains$LO_log~WildBrains$RCBR_log*WildBrains$Subspecies, robust = T)
```

```
sma(WildBrains$LOP_log~WildBrains$RCBR_log*WildBrains$Subspecies, robust = T)
sma(WildBrains$vLO_log~WildBrains$RCBR_log*WildBrains$Subspecies, robust = T)
sma(WildBrains$AME_log~WildBrains$RCBR_log*WildBrains$Subspecies, robust = T)
sma(WildBrains$AOTu_log~WildBrains$RCBR_log*WildBrains$Subspecies, robust = T)
sma(WildBrains$AL_log~WildBrains$RCBR_log*WildBrains$Subspecies, robust = T)
sma(WildBrains$LA_log~WildBrains$RCBR_log*WildBrains$Subspecies, robust = T)
sma(WildBrains$POTu_log~WildBrains$RCBR_log*WildBrains$Subspecies, robust = T)
sma(WildBrains$PB_log~WildBrains$RCBR_log*WildBrains$Subspecies, robust = T)
sma(WildBrains$MB_log~WildBrains$RCBR_log*WildBrains$Subspecies, robust = T)
sma(WildBrains$CC_log~WildBrains$RCBR_log*WildBrains$Subspecies, robust = T)


#Testing for alpha (α) changes between subspecies (elevation shifts were detected for all the neuropils but the ventral lobula, la accessory medulla, the lamina, the anterior and the posterior optic tubercle)

sma(WildBrains$ME_log~WildBrains$RCBR_log+WildBrains$Subspecies, robust = T)
sma(WildBrains$LO_log~WildBrains$RCBR_log+WildBrains$Subspecies, robust = T)
sma(WildBrains$LOP_log~WildBrains$RCBR_log+WildBrains$Subspecies, robust = T)
sma(WildBrains$vLO_log~WildBrains$RCBR_log+WildBrains$Subspecies, robust = T)
sma(WildBrains$AME_log~WildBrains$RCBR_log+WildBrains$Subspecies, robust = T)
sma(WildBrains$AOTu_log~WildBrains$RCBR_log+WildBrains$Subspecies, robust = T)
sma(WildBrains$AL_log~WildBrains$RCBR_log+WildBrains$Subspecies, robust = T)
sma(WildBrains$LA_log~WildBrains$RCBR_log+WildBrains$Subspecies, robust = T)
sma(WildBrains$POTu_log~WildBrains$RCBR_log+WildBrains$Subspecies, robust = T)
sma(WildBrains$PB_log~WildBrains$RCBR_log+WildBrains$Subspecies, robust = T)
sma(WildBrains$MB_log~WildBrains$RCBR_log+WildBrains$Subspecies, robust = T)
sma(WildBrains$CC_log~WildBrains$RCBR_log+WildBrains$Subspecies, robust =
```

T)

#Major axis shifts for neuropils sharing the same elevation in both subspecies (no detected in any case)

```
sma(WildBrains$vLO_log~WildBrains$RCBR_log + WildBrains$Subspecies, type = "shift", robust = T)
sma(WildBrains$AME_log~WildBrains$RCBR_log + WildBrains$Subspecies, type = "shift", robust = T)
sma(WildBrains$AOTu_log~WildBrains$RCBR_log + WildBrains$Subspecies, type = "shift", robust = T)
sma(WildBrains$LA_log~WildBrains$RCBR_log + WildBrains$Subspecies, type = "shift", robust = T)
sma(WildBrains$POTu_log~WildBrains$RCBR_log + WildBrains$Subspecies, type = "shift", robust = T)
```

#SMATR on wild females grouped in subspecies

#Testing for  correlation between the allometric control and the neuropil of interest (this correlation is lacking in the accessory medulla, the accesory optic tubercule and the lamina)

```
sma(WildFemales$ME_log~WildFemales$RCBR_log, robust = T)
sma(WildFemales$LO_log~WildFemales$RCBR_log, robust = T)
sma(WildFemales$LOP_log~WildFemales$RCBR_log, robust = T)
sma(WildFemales$vLO_log~WildFemales$RCBR_log, robust = T)
sma(WildFemales$AME_log~WildFemales$RCBR_log, robust = T)
sma(WildFemales$AOTu_log~WildFemales$RCBR_log, robust = T)
sma(WildFemales$AL_log~WildFemales$RCBR_log, robust = T)
sma(WildFemales$LA_log~WildFemales$RCBR_log, robust = T)
```

#Testing for slope (β) changes between subspecies (no slope differences were detected)

```
sma(WildFemales$ME_log~WildFemales$RCBR_log*WildFemales$Subspecies, robust = T)
sma(WildFemales$LO_log~WildFemales$RCBR_log*WildFemales$Subspecies, robust = T)
sma(WildFemales$LOP_log~WildFemales$RCBR_log*WildFemales$Subspecies, robust = T)
sma(WildFemales$vLO_log~WildFemales$RCBR_log*WildFemales$Subspecies, robust = T)
sma(WildFemales$AL_log~WildFemales$RCBR_log*WildFemales$Subspecies, robust = T)

sma(WildFemales$AME_log~WildFemales$RCBR_log*WildFemales$Subspecies, robust = T)
```

```
sma(WildFemales$AOTu_log~WildFemales$RCBR_log*WildFemales$Subspecies,
robust = T)
sma(WildFemales$LA_log~WildFemales$RCBR_log*WildFemales$Subspecies,
robust = T)



#Testing for alpha (α) changes between subspecies (elevation shifts were detected
for the lobula plate)

sma(WildFemales$ME_log~WildFemales$RCBR_log+WildFemales$Subspecies,
robust = T)
sma(WildFemales$LO_log~WildFemales$RCBR_log+WildFemales$Subspecies,
robust = T)
sma(WildFemales$LOP_log~WildFemales$RCBR_log+WildFemales$Subspecies,
robust = T)
sma(WildFemales$vLO_log~WildFemales$RCBR_log+WildFemales$Subspecies,
robust = T)
sma(WildFemales$AL_log~WildFemales$RCBR_log+WildFemales$Subspecies,
robust = T)
sma(WildFemales$AME_log~WildFemales$RCBR_log+WildFemales$Subspecies,
robust = T)
sma(WildFemales$AOTu_log~WildFemales$RCBR_log+WildFemales$Subspecies,
robust = T)
sma(WildFemales$LA_log~WildFemales$RCBR_log+WildFemales$Subspecies,
robust = T)


#Major axis shifts for neuropils sharing the same elevation beteen subspecies (no
detected in any case)

sma(WildBrains$ME_log~WildBrains$RCBR_log + WildBrains$Subspecies, type =
"shift", robust = T)
sma(WildBrains$LO_log~WildBrains$RCBR_log + WildBrains$Subspecies, type =
"shift", robust = T)
sma(WildBrains$vLO_log~WildBrains$RCBR_log + WildBrains$Subspecies, type =
"shift", robust = T)
sma(WildBrains$AL_log~WildBrains$RCBR_log + WildBrains$Subspecies, type =
"shift", robust = T)
sma(WildBrains$AME_log~WildBrains$RCBR_log + WildBrains$Subspecies, type =
"shift", robust = T)
sma(WildBrains$AOTu_log~WildBrains$RCBR_log + WildBrains$Subspecies, type
= "shift", robust = T)
sma(WildBrains$LA_log~WildBrains$RCBR_log + WildBrains$Subspecies, type =
"shift", robust = T)


#SMATR on wild males grouped in species
```

#Testing for  correlation between the allometric control and the neuropil of interest (this correlation is lacking in the lamina)

```
sma(WildMales$ME_log~WildMales$RCBR_log, robust = T)
sma(WildMales$LO_log~WildMales$RCBR_log, robust = T)
sma(WildMales$LOP_log~WildMales$RCBR_log, robust = T)
sma(WildMales$vLO_log~WildMales$RCBR_log, robust = T)
sma(WildMales$AME_log~WildMales$RCBR_log, robust = T)
sma(WildMales$AOTu_log~WildMales$RCBR_log, robust = T)
sma(WildMales$AL_log~WildMales$RCBR_log, robust = T)
sma(WildMales$LA_log~WildMales$RCBR_log, robust = T)
```

#Testing for slope (β) changes between subspecies (in all the cases, the slopes are similar between subspecies)

```
sma(WildMales$ME_log~WildMales$RCBR_log*WildMales$Subspecies, robust = T)
sma(WildMales$LO_log~WildMales$RCBR_log*WildMales$Subspecies, robust = T)
sma(WildMales$LOP_log~WildMales$RCBR_log*WildMales$Subspecies, robust =
T)
sma(WildMales$vLO_log~WildMales$RCBR_log*WildMales$Subspecies, robust =
T)
sma(WildMales$AME_log~WildMales$RCBR_log*WildMales$Subspecies, robust =
T)
sma(WildMales$AOTu_log~WildMales$RCBR_log*WildMales$Subspecies, robust =
T)
sma(WildMales$AL_log~WildMales$RCBR_log*WildMales$Subspecies, robust = T)

sma(WildMales$LA_log~WildMales$RCBR_log*WildMales$Subspecies, robust = T)
```

#Testing for alpha (α) changes between subspecies (elevation shifts were detected in the medula)

```
sma(WildMales$ME_log~WildMales$RCBR_log+WildMales$Subspecies, robust = T)
sma(WildMales$LO_log~WildMales$RCBR_log+WildMales$Subspecies, robust = T)
sma(WildMales$LOP_log~WildMales$RCBR_log+WildMales$Subspecies, robust =
T)
sma(WildMales$vLO_log~WildMales$RCBR_log+WildMales$Subspecies, robust =
T)
sma(WildMales$AME_log~WildMales$RCBR_log+WildMales$Subspecies, robust =
T)
sma(WildMales$AOTu_log~WildMales$RCBR_log+WildMales$Subspecies, robust =
T)
sma(WildMales$AL_log~WildMales$RCBR_log+WildMales$Subspecies, robust = T)
sma(WildMales$LA_log~WildMales$RCBR_log+WildMales$Subspecies, robust = T)
```

#Major axis shifts for neuropils sharing the same elevation between subspecies (no detected in any case)

```
sma(WildBrains$LO_log~WildBrains$RCBR_log + WildBrains$Subspecies, type = "shift", robust = T)
sma(WildBrains$LOP_log~WildBrains$RCBR_log + WildBrains$Subspecies, type = "shift", robust = T)
sma(WildBrains$vLO_log~WildBrains$RCBR_log + WildBrains$Subspecies, type = "shift", robust = T)
sma(WildBrains$AME_log~WildBrains$RCBR_log + WildBrains$Subspecies, type = "shift", robust = T)
sma(WildBrains$AOTu_log~WildBrains$RCBR_log + WildBrains$Subspecies, type = "shift", robust = T)
sma(WildBrains$AL_log~WildBrains$RCBR_log + WildBrains$Subspecies, type = "shift", robust = T)


sma(WildBrains$LA_log~WildBrains$RCBR_log + WildBrains$Subspecies, type = "shift", robust = T)


#SMATR on wild erato individuals as a whole grouped in sexes (no correlation tests needed as these were performed earlier)

#Testing for slope (β) changes between sexes (in all the neuropiles, the slopes are similar)

sma(WildBrains$ME_log~WildBrains$RCBR_log*WildBrains$Sex, robust = T)
sma(WildBrains$LO_log~WildBrains$RCBR_log*WildBrains$Sex, robust = T)
sma(WildBrains$LOP_log~WildBrains$RCBR_log*WildBrains$Sex, robust = T)
sma(WildBrains$vLO_log~WildBrains$RCBR_log*WildBrains$Sex, robust = T)
sma(WildBrains$AME_log~WildBrains$RCBR_log*WildBrains$Sex, robust = T)
sma(WildBrains$AOTu_log~WildBrains$RCBR_log*WildBrains$Sex, robust = T)
sma(WildBrains$AL_log~WildBrains$RCBR_log*WildBrains$Sex, robust = T)
sma(WildBrains$LA_log~WildBrains$RCBR_log*WildBrains$Sex, robust = T)

#Testing for alpha (α) changes between sexes (elevation shifts were detected for the medulla, the lobula, the lobula plate, the ventral lobula, the antennal lobe and the lamina)

sma(WildBrains$ME_log~WildBrains$RCBR_log+WildBrains$Sex, robust = T)
sma(WildBrains$LO_log~WildBrains$RCBR_log+WildBrains$Sex, robust = T)
sma(WildBrains$LOP_log~WildBrains$RCBR_log+WildBrains$Sex, robust = T)
sma(WildBrains$vLO_log~WildBrains$RCBR_log+WildBrains$Sex, robust = T)
sma(WildBrains$AME_log~WildBrains$RCBR_log+WildBrains$Sex, robust = T)
sma(WildBrains$AOTu_log~WildBrains$RCBR_log+WildBrains$Sex, robust = T)
sma(WildBrains$AL_log~WildBrains$RCBR_log+WildBrains$Sex, robust = T)
sma(WildBrains$LA_log~WildBrains$RCBR_log+WildBrains$Sex, robust = T)


#Major axis shifts for neuropils sharing the same elevation in both subspecies (no
```

shifts detected)

```
sma(WildBrains$AME_log~WildBrains$RCBR_log + WildBrains$Sex, type = "shift", robust = T)
sma(WildBrains$AOTu_log~WildBrains$RCBR_log + WildBrains$Sex, type = "shift", robust = T)
```

#SMATR on wild chestertonii individuals grouped in sexes

#Testing for correlation between the allometric control and the neuropil of interest within wild chestertonii (this is lacking for the accessory medulla and the lamina)

```
sma(Wildchestertonii$ME_log~Wildchestertonii$RCBR_log, robust = T)
sma(Wildchestertonii$LO_log~Wildchestertonii$RCBR_log, robust = T)
sma(Wildchestertonii$LOP_log~Wildchestertonii$RCBR_log, robust = T)
sma(Wildchestertonii$vLO_log~Wildchestertonii$RCBR_log, robust = T)
sma(Wildchestertonii$AME_log~Wildchestertonii$RCBR_log, robust = T)
sma(Wildchestertonii$AOTu_log~Wildchestertonii$RCBR_log, robust = T)
sma(Wildchestertonii$AL_log~Wildchestertonii$RCBR_log, robust = T)
sma(Wildchestertonii$LA_log~Wildchestertonii$RCBR_log, robust = T)
```

#Testing for slope (β) changes between sexes within chestertonii (in all the neuropiles, the slopes are similar)

```
sma(Wildchestertonii$ME_log~Wildchestertonii$RCBR_log*Wildchestertonii$Sex, robust = T)
sma(Wildchestertonii$LO_log~Wildchestertonii$RCBR_log*Wildchestertonii$Sex, robust = T)
sma(Wildchestertonii$LOP_log~Wildchestertonii$RCBR_log*Wildchestertonii$Sex, robust = T)
sma(Wildchestertonii$vLO_log~Wildchestertonii$RCBR_log*Wildchestertonii$Sex, robust = T)
sma(Wildchestertonii$AOTu_log~Wildchestertonii$RCBR_log*Wildchestertonii$Sex, robust = T)
sma(Wildchestertonii$AL_log~Wildchestertonii$RCBR_log*Wildchestertonii$Sex, robust = T)
sma(Wildchestertonii$AME_log~Wildchestertonii$RCBR_log*Wildchestertonii$Sex, robust = T)
sma(Wildchestertonii$LA_log~Wildchestertonii$RCBR_log*Wildchestertonii$Sex, robust = T)
```

#Testing for alpha (α) changes between sexes within chestertonii (elevation shifts were detected for the lobula and the lobula plate)

```
sma(Wildchestertonii$ME_log~Wildchestertonii$RCBR_log+Wildchestertonii$Sex, robust = T)
```

```
sma(Wildchestertonii$LO_log~Wildchestertonii$RCBR_log+Wildchestertonii$Sex, robust = T)
sma(Wildchestertonii$LOP_log~Wildchestertonii$RCBR_log+Wildchestertonii$Sex, robust = T)
sma(Wildchestertonii$vLO_log~Wildchestertonii$RCBR_log+Wildchestertonii$Sex, robust = T)
sma(Wildchestertonii$AOTu_log~Wildchestertonii$RCBR_log+Wildchestertonii$Sex, robust = T)
sma(Wildchestertonii$AL_log~Wildchestertonii$RCBR_log+Wildchestertonii$Sex, robust = T)
sma(Wildchestertonii$AME_log~Wildchestertonii$RCBR_log+Wildchestertonii$Sex, robust = T)
sma(Wildchestertonii$LA_log~Wildchestertonii$RCBR_log+Wildchestertonii$Sex, robust = T)

#Major axis shifts for neuropils sharing the same elevation between sexes within chestertonii (no shifts detected)

sma(Wildchestertonii$ME_log~Wildchestertonii$RCBR_log + Wildchestertonii$Sex, type = "shift", robust = T)
sma(Wildchestertonii$AOTu_log~Wildchestertonii$RCBR_log + Wildchestertonii$Sex, type = "shift", robust = T)
sma(Wildchestertonii$AL_log~Wildchestertonii$RCBR_log + Wildchestertonii$Sex, type = "shift", robust = T)
sma(Wildchestertonii$AME_log~Wildchestertonii$RCBR_log + Wildchestertonii$Sex, type = "shift", robust = T)
sma(Wildchestertonii$LA_log~Wildchestertonii$RCBR_log + Wildchestertonii$Sex, type = "shift", robust = T)


#SMATR on wild venus individuals grouped in sexes

#Testing for correlation between the allometric control and the neuropil of interest within wild venus (this is present for all the neuropiles)

sma(Wildvenus$ME_log~Wildvenus$RCBR_log, robust = T)
sma(Wildvenus$LO_log~Wildvenus$RCBR_log, robust = T)
sma(Wildvenus$LOP_log~Wildvenus$RCBR_log, robust = T)
sma(Wildvenus$vLO_log~Wildvenus$RCBR_log, robust = T)
sma(Wildvenus$AME_log~Wildvenus$RCBR_log, robust = T)
sma(Wildvenus$AOTu_log~Wildvenus$RCBR_log, robust = T)
sma(Wildvenus$AL_log~Wildvenus$RCBR_log, robust = T)
sma(Wildvenus$LA_log~Wildvenus$RCBR_log, robust = T)


#Testing for slope (β) changes between sexes within venus (in all the neuropiles, the slopes are similar)

sma(Wildvenus$ME_log~Wildvenus$RCBR_log*Wildvenus$Sex, robust = T)
sma(Wildvenus$LO_log~Wildvenus$RCBR_log*Wildvenus$Sex, robust = T)
```

```
sma(Wildvenus$LOP_log~Wildvenus$RCBR_log*Wildvenus$Sex, robust = T)
sma(Wildvenus$vLO_log~Wildvenus$RCBR_log*Wildvenus$Sex, robust = T)
sma(Wildvenus$AME_log~Wildvenus$RCBR_log*Wildvenus$Sex, robust = T)
sma(Wildvenus$AOTu_log~Wildvenus$RCBR_log*Wildvenus$Sex, robust = T)
sma(Wildvenus$AL_log~Wildvenus$RCBR_log*Wildvenus$Sex, robust = T)
sma(Wildvenus$LA_log~Wildvenus$RCBR_log*Wildvenus$Sex, robust = T)

#Testing for alpha (α) changes between sexes within venus (elevation shifts were detected for the medulla, the antennal lobe and the lamina)

sma(Wildvenus$ME_log~Wildvenus$RCBR_log+Wildvenus$Sex, robust = T)
sma(Wildvenus$LO_log~Wildvenus$RCBR_log+Wildvenus$Sex, robust = T)
sma(Wildvenus$LOP_log~Wildvenus$RCBR_log+Wildvenus$Sex, robust = T)
sma(Wildvenus$vLO_log~Wildvenus$RCBR_log+Wildvenus$Sex, robust = T)
sma(Wildvenus$AME_log~Wildvenus$RCBR_log+Wildvenus$Sex, robust = T)
sma(Wildvenus$AOTu_log~Wildvenus$RCBR_log+Wildvenus$Sex, robust = T)
sma(Wildvenus$AL_log~Wildvenus$RCBR_log+Wildvenus$Sex, robust = T)
sma(Wildvenus$LA_log~Wildvenus$RCBR_log+Wildvenus$Sex, robust = T)

#Major axis shifts for neuropils sharing the same elevation between sexes within venus (no shifts detected)

sma(Wildvenus$LO_log~Wildvenus$RCBR_log + Wildvenus$Sex, type = "shift", robust = T)
sma(Wildvenus$LOP_log~Wildvenus$RCBR_log + Wildvenus$Sex, type = "shift", robust = T)
sma(Wildvenus$vLO_log~Wildvenus$RCBR_log + Wildvenus$Sex, type = "shift", robust = T)
sma(Wildvenus$AME_log~Wildvenus$RCBR_log + Wildvenus$Sex, type = "shift", robust = T)
sma(Wildvenus$AOTu_log~Wildvenus$RCBR_log + Wildvenus$Sex, type = "shift", robust = T)


#Parallelism in brain divergence analyses with linear mixed models

#Importing the data

Brainmix

#Models on neuropil size across localities, "*" means that effect is signifiant

#Modelling the effects of habitat*, locality* and their interaction on ME size

MEcombinedfull<-lmer(ME_log ~ RCBR_log+Habitat*Locality+(1|Sex), data=Brainmix)

drop1(MEcombinedfull, test="Chisq")
newMEcombined<-update(MEcombinedfull, ~.-Habitat:Locality)
```

```
drop1(newMEcombined, test="Chisq")

MEcombinedreduced<-lmer(ME_log ~ RCBR_log+Habitat+Locality+(1|Sex),
data=Brainmix)
MEcombinedalt<-lmer(ME_log ~RCBR_log+Locality+(1|Sex), data=Brainmix)

anova(MEcombinedreduced, MEcombinedalt)

Anova(MEcombinedreduced)

summary(MEcombinedreduced)

r2beta(MEcombinedreduced, method = "kr")

r2beta(model=MEcombinedreduced,partial=TRUE,method='kr')
plot(x=r2)
print(r2)


#Modelling the effects of habitat*, locality* and their interaction on LOP size

LOPcombinedfull<-lmer(LOP_log ~ RCBR_log+Habitat*Locality+(1|Sex),
data=Brainmix)

drop1(LOPcombinedfull, test="Chisq")
newLOPcombined<-update(LOPcombinedfull, ~.-Habitat:Locality)

drop1(newLOPcombined, test="Chisq")

LOPcombinedreduced<-lmer(LOP_log ~ RCBR_log+Habitat+Locality+(1|Sex),
data=Brainmix)
LOPcombinedalt<-lmer(LOP_log ~RCBR_log+Locality+(1|Sex), data=Brainmix)

anova(LOPcombinedreduced, LOPcombinedalt)

Anova(LOPcombinedreduced)

summary(LOPcombinedreduced)


r2beta(LOPcombinedreduced, method = "kr")

r2beta(model=LOPcombinedreduced,partial=TRUE,method='kr')
plot(x=r2)
print(r2)


#Modelling the effects of habitat*, locality* and their interaction on LO size

LOcombinedfull<-lmer(LO_log ~ RCBR_log+Habitat*Locality+(1|Sex),
```

```
data=Brainmix)

drop1(LOcombinedfull, test="Chisq")
newLOcombined<-update(LOcombinedfull, ~.-Habitat:Locality)

drop1(newLOcombined, test="Chisq")

LOcombinedreduced<-lmer(LO_log ~ RCBR_log+Habitat+Locality+(1|Sex),
data=Brainmix)
LOcombinedalt<-lmer(LO_log ~RCBR_log+Locality+(1|Sex), data=Brainmix)

anova(LOcombinedreduced, LOcombinedalt)

Anova(LOcombinedreduced)

summary(LOcombinedreduced)

r.squaredGLMM(LOcombinedreduced)

r2beta(LOcombinedreduced, method = "kr")

r2beta(model=LOcombinedreduced,partial=TRUE,method='kr')
plot(x=r2)
print(r2)


#Modelling the effects of habitat, locality* and their interaction* on AL size

ALcombinedfull<-lmer(AL_log ~ RCBR_log+Habitat*Locality+(1|Sex),
data=Brainmix)

drop1(ALcombinedfull, test="Chisq")

ALcombinedreduced<-lmer(AL_log ~ RCBR_log+Habitat*Locality+(1|Sex),
data=Brainmix)
ALcombinedalt<-lmer(AL_log ~RCBR_log+Habitat+Locality+(1|Sex), data=Brainmix)

anova(ALcombinedreduced, ALcombinedalt)

Anova(ALcombinedreduced)

summary(ALcombinedreduced)

r2beta(ALcombinedreduced, method = "sgv")

r2 = r2beta(model=ALcombinedreduced,partial=TRUE,method='sgv')
plot(x=r2)
print(r2)
```
